# Defective Schwann cell lipid metabolism alters plasma membrane dynamics in Charcot-Marie-Tooth disease 1A

**DOI:** 10.1101/2023.04.02.535224

**Authors:** Robert Prior, Alessio Silva, Tim Vangansewinkel, Jakub Idkowiak, Arun Kumar Tharkeshwar, Tom P. Hellings, Iliana Michailidou, Jeroen Vreijling, Maarten Loos, Bastijn Koopmans, Nina Vlek, Nina Straat, Cedrick Agaser, Tom Kuipers, Christine Michiels, Elisabeth Rossaert, Stijn Verschoren, Wendy Vermeire, Vincent de Laat, Jonas Dehairs, Kristel Eggermont, Diede van den Biggelaar, Adekunle T. Bademosi, Frederic A. Meunier, Martin vandeVen, Philip Van Damme, Hailiang Mei, Johannes V. Swinnen, Ivo Lambrichts, Frank Baas, Kees Fluiter, Esther Wolfs, Ludo Van Den Bosch

## Abstract

Duplication of *PMP22* causes Charcot-Marie-Tooth disease type 1A (CMT1A) and is known to disrupt the lipid metabolism in myelinating Schwann cells by unknown mechanisms. By using two CMT1A mouse models overexpressing human *PMP22*, we discovered that *PMP22* dose-dependently downregulates genes that are involved in lipid and cholesterol metabolism. Lipidomic analysis on CMT1A mouse sciatic nerves confirmed lipid metabolic abnormalities primarily associated with cholesterol and sphingolipids. We observed similar lipidomic profiles and downregulation of genes associated with lipid metabolism in human CMT1A patient induced pluripotent stem cell-derived Schwann cell precursors (iPSC-SCPs). We confirmed these findings by demonstrating altered lipid raft dynamics and plasma membrane fluidity in CMT1A iPSC-SCPs. Additionally, we identified impaired cholesterol incorporation in the plasma membrane due to altered lipid storage homeostasis in CMT1A iPSC-SCPs, which could be modulated by changing the lipid composition of the cell culture medium. These findings suggest that PMP22 plays a role in regulating the lipid composition of the plasma membrane and lipid storage homeostasis. Targeting lipid metabolism may hold promise as a potential treatment for CMT1A patients.

**Graphical abstract:** 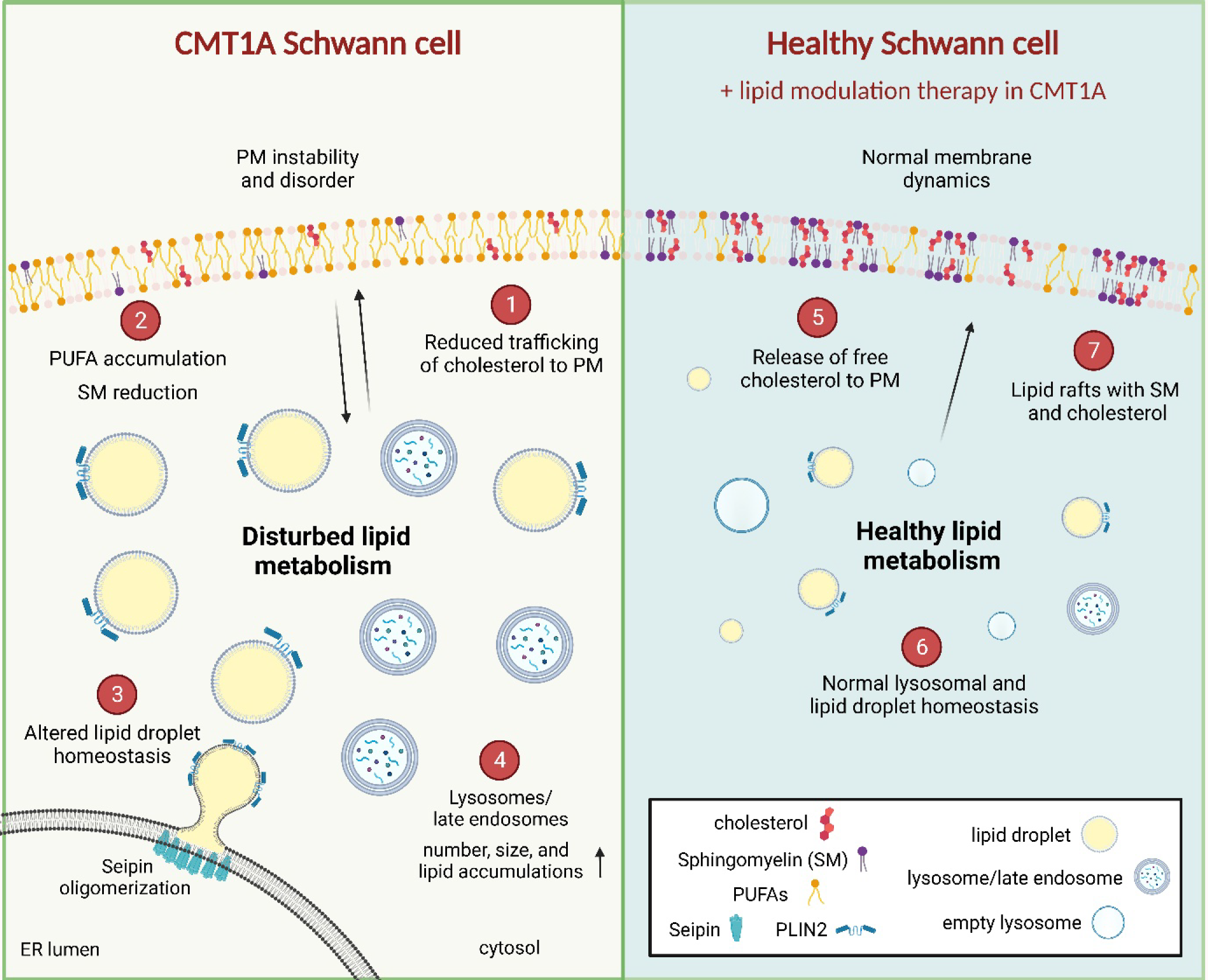

**Highlights:**

- *PMP22* copy number causes a dose-dependent suppression of cholesterol and lipid biosynthesis in peripheral nerves of CMT1A mice
- Lipid composition is altered in the sciatic nerves of CMT1A mice and in the membranes of patient derived iPSC-SCPs, with a significant reduction in sphingolipids
- CMT1A iPSC-SCPs show decreased plasma membrane lipids required for regulating lipid raft dynamics, membrane fluidity, and membrane order
- Lipid storage misregulation is key in the pathogenesis of CMT1A

## Introduction

Duplications of the *PMP22* gene, which encodes peripheral myelin protein 22 (PMP22), give rise to the most prevalent form of Charcot-Marie-Tooth disease (CMT), CMT1A (Raeymaekers et al., 1991; Timmerman et al., 1992). CMT1A patients exhibit a dysmyelinating phenotype in their peripheral nerves and may experience mild-to-severe muscular atrophy in the distal regions of their body, with some also presenting with sensory symptoms (van Paassen et al., 2014). PMP22 is a membrane glycoprotein constituting 2 to 5% of compact myelin of the peripheral nervous system (PNS), which is produced by Schwann cells (Amici et al., 2007).

Myelin is a complex and intricate structure that comprises both compacted and non-compacted membranes, and is not only essential for the maturation of associated axons but also to increase the speed of electrical nerve conduction (Brady et al., 1999; Salzer and Zalc, 2016). Myelin consists out of about 70 to 80% lipids, which are mainly fatty aldehydes and very long-chain fatty acids. The remaining 20 to 30% are proteins and over 60% of these are glycoproteins (Liu et al., 2019; Martini, 2001; Siems et al., 2020). Cholesterol is a critical component of the myelin membrane as it influences myelination in several ways. It is a vital component that influences glial cell differentiation, myelin membrane biogenesis, and the formation of functional myelin (Pertusa et al., 2007).

In the plasma membrane (PM), cholesterol is key in the regulation of membrane fluidity. It can partition into more ordered and stable phases than the surrounding fluidic phase, where it is enriched together with a specific type of sphingolipid (SP), called sphingomyelin (SM). These liquid-ordered domains are called lipid rafts and are specialized regions of the PM that contain proteins with different functions, including the shuttling of molecules on the cell surface and the organization of cell signal transduction pathways (Lingwood and Simons, 2010; Simons and Ehehalt, 2002; Simons and Toomre, 2000).

Importantly, lipid rafts provide a platform for the clustering of multiple molecules, such as lipids and receptors, allowing for efficient and specific signaling during myelination. Here, cholesterol helps to maintain the fluidity and stability of the PM (Crane and Tamm, 2004; Redondo-Morata et al., 2012). Cholesterol trafficking and the establishment of lipid rafts on the cell membrane is impaired in *Pmp22*-deficient mice (Lee et al., 2014; Zhou et al., 2019). However, it is unknown whether CMT1A patient cells have an altered lipid metabolism or impaired lipid raft function.

The storage of cholesterol and lipids in eukaryotic cells is via specialized organelles called lipid droplets (LDs). LDs can act as an endogenous source of lipids for biomembranes. From here, fatty acids can be mobilized by the cell during nutrient stress, cell differentiation, or biomembrane repair via lipophagy (i.e., the lysosome-mediated recycling or breakdown of LDs) or the enzyme-mediated lipolysis, which is primarily executed via adipose triglyceride lipase (ATGL) [reviewed in (Olzmann and Carvalho, 2019)] (Rambold et al., 2015; Sztalryd and Brasaemle, 2017). During lipid starvation, cells initially produce LDs as a short-term cellular response, followed by their breakdown and use when starvation is prolonged [reviewed in (Henne et al., 2018)].

Currently, *PMP22* overexpressing mouse models, such as the C22 and the C3 CMT1A mouse models, have helped unravel some of the pathogenic mechanism underlying CMT1A aetiology. These CMT1A mouse models overexpress different copy number levels of human *PMP22*, giving rise to varying disease severity between the two mouse models. The disease progression in the C22 mouse model has been described previously (Chittoor et al., 2013; Fortun et al., 2006; Huxley et al., 1996; Robaglia-Schlupp et al., 2002; Verhamme et al., 2011), and is considered as a severe dysmyelinating phenotype. Harboring at least 7 copies of the yeast artificial chromosome containing the human *PMP22* gene, the C22 mice display postnatal defects in myelination (Robaglia-Schlupp *et al*., 2002; Robertson et al., 1999; Verhamme *et al*., 2011). The spontaneous revertant of the C22 mouse, designated the C3 mouse model (Verhamme *et al*., 2011) contains 5 copies of the human *PMP22* gene, as we reported previously (Prior et al., 2022). These mouse models, in addition to the transgenic CMT1A rat model (Fledrich et al., 2012) are the current gold standard *in vivo* models for CMT1A research. Unfortunately, all transgenic models have their inherent flaws and can never fully recapitulate the complexity of human diseases. However, the development of human induced pluripotent stem cells (iPSCs) has rapidly facilitated the study of patient derived cells, enabling researchers to gain further insight into human diseases.

In this study, we aimed at understanding how CMT1A mice and patient-derived cells handle lipids during Schwann cell development. In particular, we used human iPSCs to model CMT1A by differentiating them into Schwann cell precursors (iPSC-SCPs). We found that increased *PMP22* copy number alters the lipid flux in both CMT1A mice and patient derived iPSC-SCPs. This alteration in the lipidome was characterized by alterations in lipid storage homeostasis, lipid recycling and PM lipid composition. Moreover, this change in lipid composition led to disordered PM lipid properties and lipid raft dynamics. Ultimately, we showed that by targeting lipid metabolic processes, we were able to restore the cholesterol and lipid recycling defects in CMT1A patient iPSC-SCPs derived cells, further highlighting lipid metabolism as a targetable therapy for CMT1A patients.

## Results

### Cholesterol biosynthesis is the major dysregulated pathway in CMT1A mice

Similar to the C22 mouse model (Verhamme et al., 2011), the C3 mice have postnatal myelination defects (Supplemental Fig. 1a). Using immunohistochemistry, the level of human PMP22 expression is prominently visible in the C22 and is subtly visible in the sciatic nerves of the C3 mouse models (Fig. 1a). To investigate the effect dose-dependent *PMP22* overexpression has on murine peripheral nerve development, we analyzed the evolution of the sciatic nerves’ transcriptome throughout development (at 3, 5, 7, 9, and 12 weeks of age) using bulk RNA-sequencing (overview in Fig. 1b). Our results indicate that cholesterol biosynthesis (Fig. 1c, d) and lipid metabolism (Fig. 1e) are the major dysregulated pathways in these CMT1A mouse models when compared to their age-matched wild-type (WT) littermates. Interestingly, cholesterol biosynthesis is progressively upregulated in C3 mice throughout development, while it is continuously repressed in C22 mice (Fig. 1c, d). This indicates that *PMP22* overexpression has a dose-dependent inhibitory effect on cholesterol and lipid metabolism.

**Figure 1.**
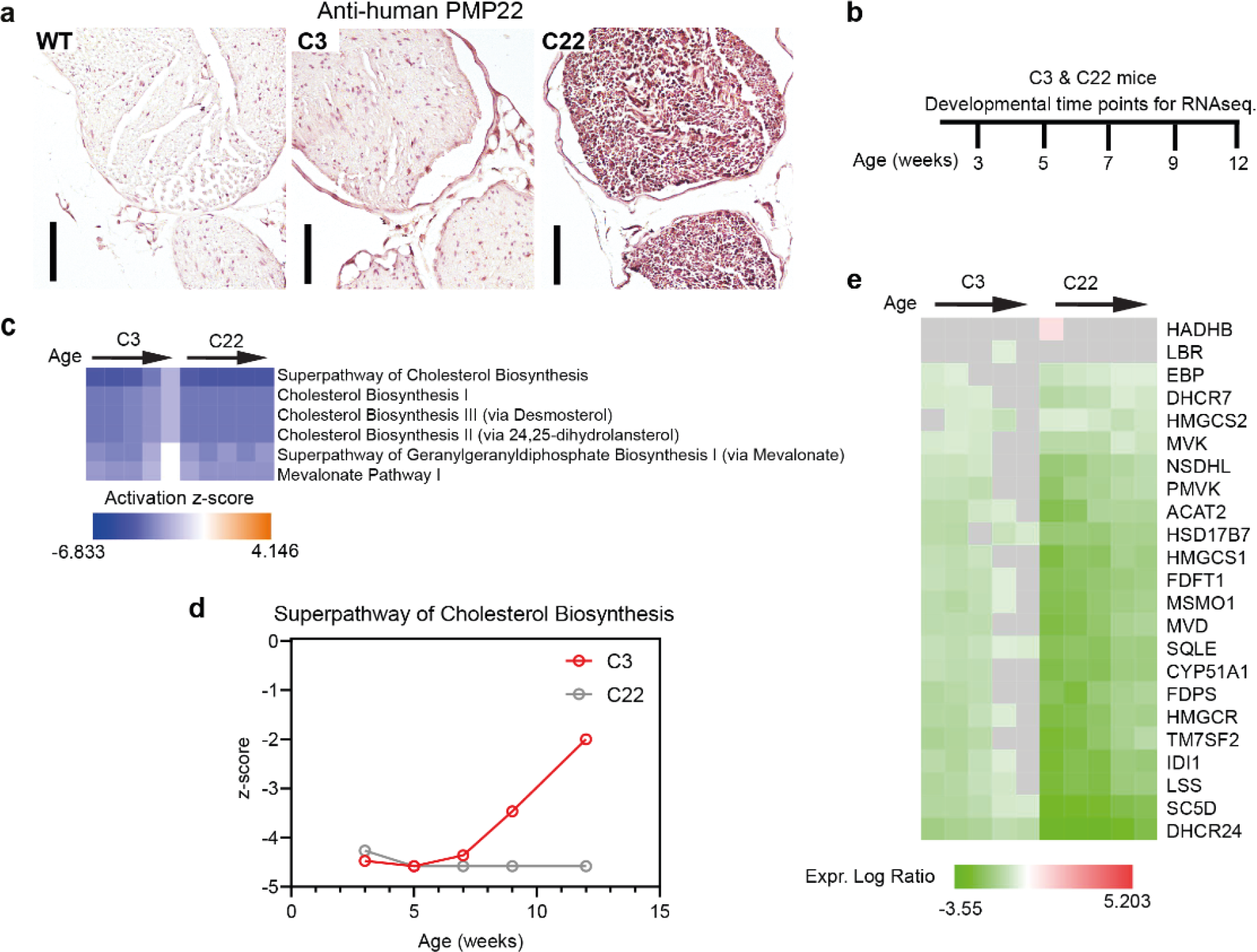
Cholesterol biosynthesis is the major dysregulated pathway during nerve development of CMT1A mice. **a)** Immunohistochemical analysis of human PMP22 in the sciatic nerves of the C3 and C22 mouse models at 12 weeks-of-age. **b)** Schematic overview of the time points analyzed for bulk RNA-sequencing of sciatic nerves isolated from the C3 and C22 mouse models with their age-matched littermate controls used for normalization. **c)** Activation heatmap of the main canonical pathways (the cholesterol biosynthesis-related pathways) that are dysregulated in both the C3 and C22 mouse models throughout their postnatal development when compared to their littermate controls. **d)** The superpathway of cholesterol biosynthesis z-scores are plotted over the developmental time points at which data collection was conducted. Littermate control z-scores are equivalent to 0 on the y-axis of the graph. **e)** Heatmap illustrating the temporal expression profiles of dysregulated lipid metabolism-related genes in the C3 and C22 mouse models throughout development. Scale bar in (a) = 100 µm. Black arrows in (c) and (e) represent developmental time points from 3-12 weeks in age, as displayed in (b).

### *PMP22* overexpression causes abnormal lipid metabolism in the C3 mouse model

To investigate the role PMP22 has in cholesterol and lipid metabolism in more detail, we focused on the less severe C3 CMT1A mouse model. These C3 mice present a dysmyelinating phenotype early in development followed by an improvement in myelination (Prior *et al*., 2022; Verhamme *et al*., 2011), similar to the aforementioned partial developmental amelioration in the cholesterol biosynthesis and lipid metabolism pathways (Fig. 1c, d). This results in thinner myelin sheaths and reduced axon caliber size in the peripheral nerves of adult mice (Supplemental Fig. 1a), as previously described and quantified (Verhamme *et al*., 2011). C3 mice developed a mild motor phenotype at three weeks of age onwards (Supplemental Fig. 1b-h). Using a time point at five weeks of age, we examined the lipid composition in the sciatic nerves of the C3 mice and compared it with that of their age-matched WT littermates. We observed a clear separation between the two mouse populations using principal component analysis (PCA) (Fig. 2a). A marked number of sphingolipids (SPs), especially sphingomyelin (SM), mono-hexosylceramide (HexCer), dihydroceramide (dhCer), and di-hexosylceramide (Hex2Cer) were significantly downregulated in C3 mice compared to WT controls (Fig. 2b). Contrary to the other SPs, most ceramide (Cer) lipids were upregulated in C3 mice, albeit without reaching statistical significance. Among the GL, the highly unsaturated TG were the most significantly downregulated. In the case of glycerophospholipids (GPL), the most significant alterations occurred in phosphatidylethanolamine (PE), PE plasmalogens (P-), PE ethers (O-), PC, PC P-, PC O-, and PS. In addition, higher levels of long chain (36, 38, 40, 42, and 44 total carbon number) and unsaturated PC species were observed, however, without reaching statistical significance. Lysoglycerophospholipids (lysophosphatidylcholine, LPC, and lysophosphatidylethanolamine, LPE), phosphatidylinositol (PI), and phosphatidylglycerol (PG) were mainly downregulated in the C3 mice but only single lipids showed significant changes (Fig. 2b, d). Moreover, cholesterol levels were confirmed to be downregulated in the sciatic nerves of the C3 mouse model at this developmental time point (Fig. 2c), in line with bulk RNA sequencing results (Fig. 1c, d). Using a simplified lipid network, the sciatic nerve lipidome of the C3 mice was compared to the one of their WT littermates (Fig. 2d). Overall, these data indicate that sciatic nerve lipid homeostasis is severely dysregulated in the C3 CMT1A mouse model.

**Figure 2.**
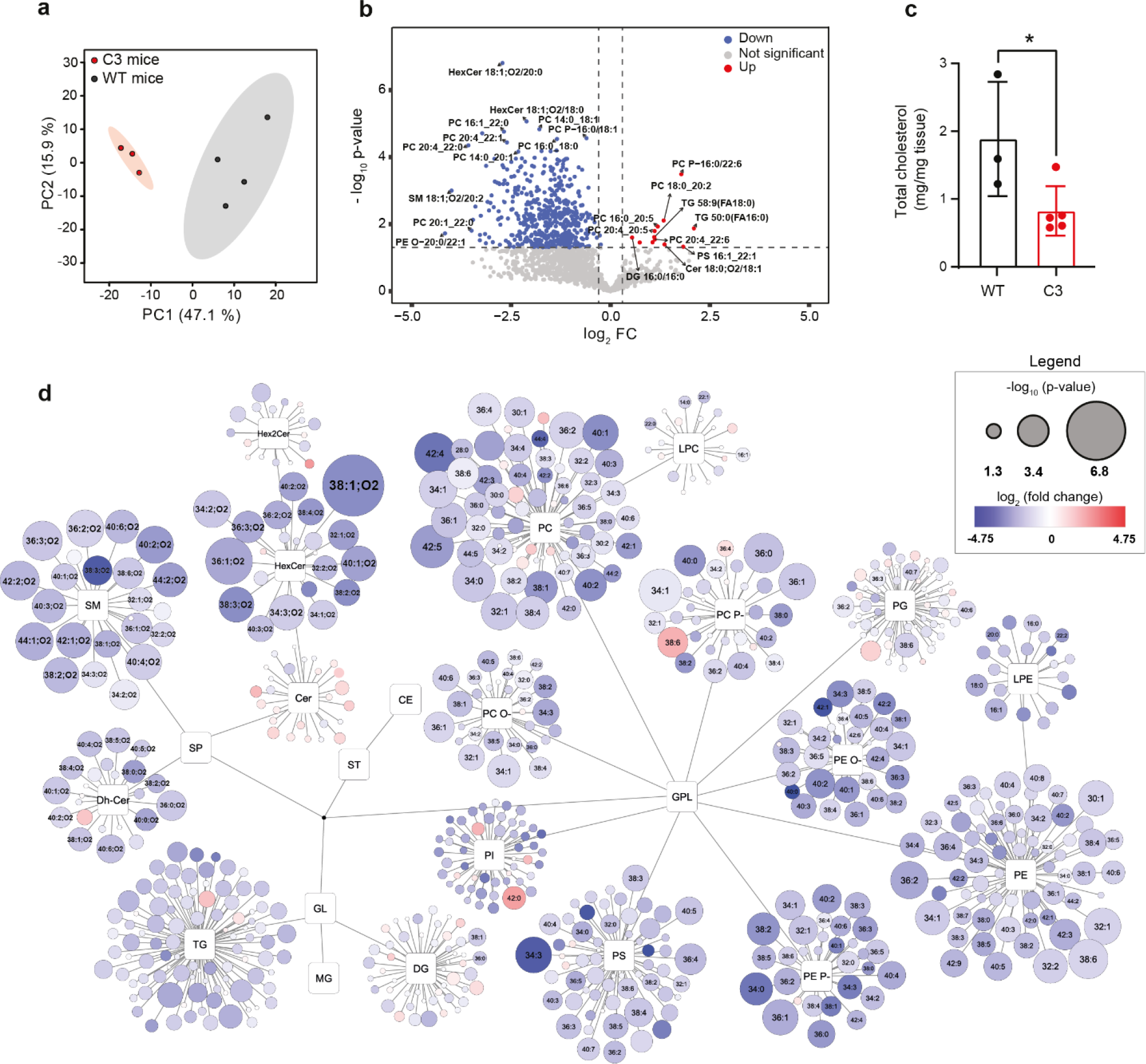
Lipidomic analysis highlights major alterations in the expression of lipid species in the sciatic nerves of 5-weeks-old C3 mice. **a)** Principal component analysis (PCA) plot of lipids in sciatic nerves from C3 mice versus their WT littermate controls at 5 weeks of age. The top principal components 1 and 2 (PC1 and PC2, respectively) were used to generate the graph. **b)** Volcano plot demonstrating the relative expression of lipid species between C3 mice and WT controls. P-value threshold was set at 0.05; Fold change (FC) threshold: log2(FC) = 0.3. **c)** Total cholesterol measurement from sciatic nerves of 5-week-old C3 mice and WT littermates. Statistical significance was evaluated using a two-tailed, unpaired t-test (* p < 0.05). Data are represented as mean ± S.D. **d)** Simplified network visualization of lipid metabolism showing alterations in lipid profiles of C3 mice compared to their WT littermates. Circles represent the detected lipid species, where the circle size expresses the significance according to p-value, while the color darkness defines the degree of up/downregulation (red/blue) according to the fold change. The most discriminating lipids are annotated. Source data are provided as a Source Data file. The number of mice used in a, b, d) was as follows: WT mice = 4, and C3 mice = 3. In c) the number of mice used was as follows: WT mice = 3, and C3 mice = 5.

### Generation of Schwann cells from induced pluripotent stem cells

To model CMT1A *in vitro*, we used a protocol modified from Kim et al. (Kim et al., 2017) to generate iPSC-derived Schwann cells (iPSC-SCs) and iPSC-SCPs (Fig. 3a). Using this protocol, we observed morphological changes from an iPSC colony state to a Schwann cell-like morphology, accompanied by the expression of key proteins relating to iPSCs, early neural crest cells, SCPs, and eventually Schwann cells (Fig. 3b; Supplemental Fig. 2 and 3). Moreover, we detected a temporal expression profile of genes essential to Schwann cell lineage development (Fig. 3c). As a demonstration of functionality, we were able to show that these iPSC-SCs produced myelin when cocultured with mouse dorsal root ganglion (DRG) neurons (Fig. 3d).

**Figure 3.**
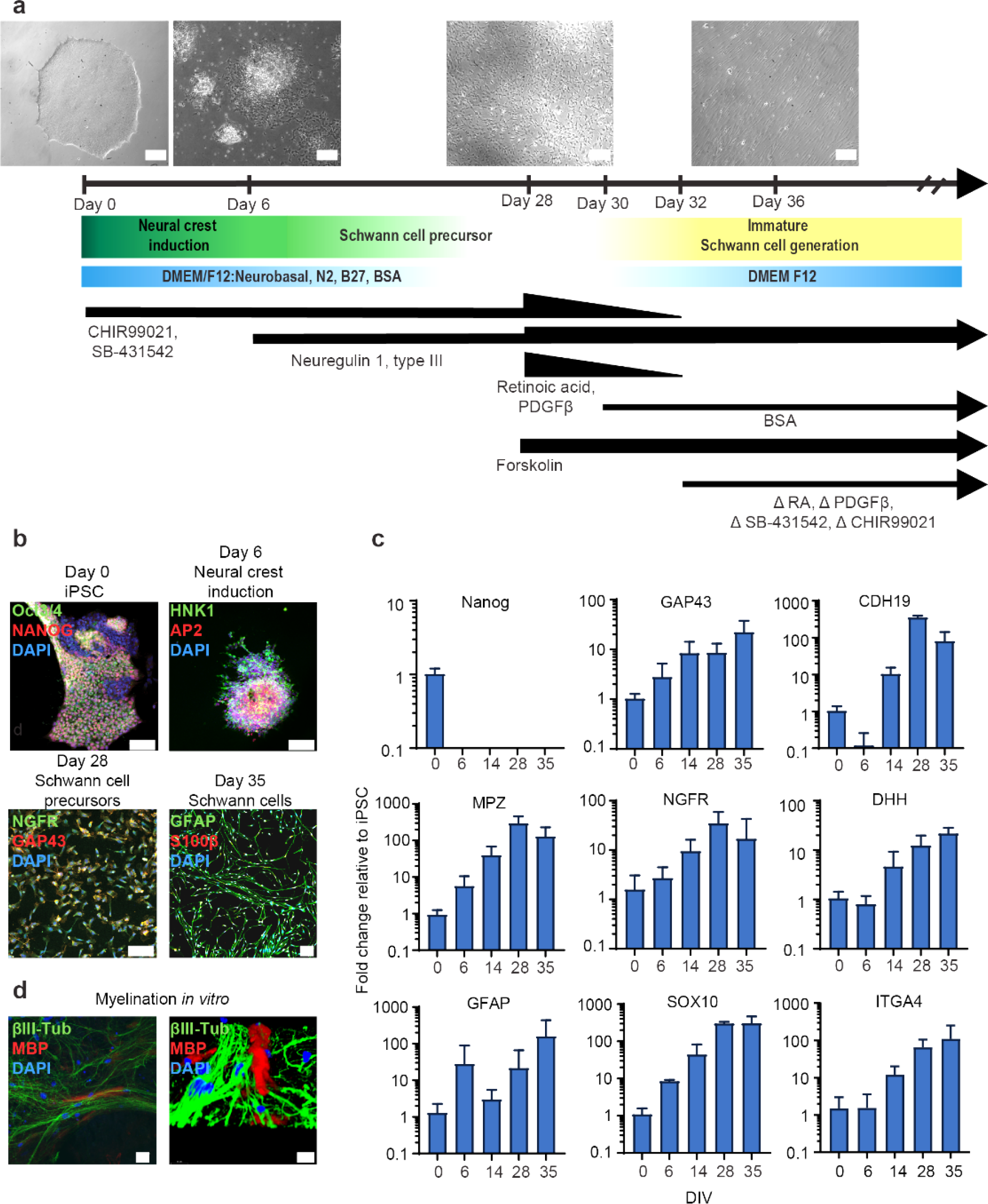
Protocol to generate Schwann cells from induced pluripotent stem cells (iPSCs) **a)** Schematic overview of the iPSC-SC protocol with key medium composition component incorporation or removal (Δ) indicated. Scale bar = 100 µm. **b)** Immunocytochemistry staining of key lineage markers representing a switch in phenotypes at days 0, 6, 28, and 35 from the protocol. Scale bar = 50 µm. **c)** qPCR temporal mRNA expression profiles of *NANOG*, *GAP43*, *CDH19*, *MPZ*, *NGFR*, *DHH*, *GFAP*, *SOX10*, and *ITGA4* quantifications. DIV: day *in vitro*. Data are represented as mean ± S.D. **d)** Immunocytochemistry staining of iPSC-SCs indicate expression of myelin basic protein (MBP) after being co-cultured with mouse DRGs for >8 weeks. Scale bar = 20 µm. The image on the right shows a 3D reconstruction of the image on the left. Scale bar = 10 µm.

### CMT1A iPSC-SCPs have dysregulated transcriptional and lipidome networks

We used a CMT1A patient line, CS67i-CMT-n1, and its isogenic control iPSC line, isogenic-CS67i-CMT (referred to as “CMT1A” and “isogenic” lines, respectively, from hereon) to generate iPSC-SCPs (Supplemental Fig. 4 and 5) to study early pathological abnormalities in CMT1A Schwann cell lineage development using bulk RNA-sequencing (Fig. 4a). The CMT1A iPSC patient line contained one extra copy of *PMP22*, while the isogenic is a genetically restored iPSC control of the CMT1A line (Supplemental Fig. 6). The transcriptome of both the CMT1A and the isogenic iPSC-SCPs is similar to that of neural crest progenitors (Supplemental Fig. 7) (Rayon et al., 2021). *PMP22* mRNA levels fluctuated in the CMT1A iPSC-SCP between overexpressed and reduced in comparison to the isogenic iPSC-SCPs depending on whether cells were cultured in high bovine serum albumin (BSA) conditions (0.5%) or low BSA conditions (0.005%), respectively (Supplemental Fig. 8; Fig. 4a). However, iPSC-SCPs were maintained in standard low BSA culture concentrations, as high BSA conditions can impair cell culture maintenance (data not shown). Several Schwann cell development and myelin-related genes, in addition to lipid metabolism and autophagy genes, were dysregulated in the CMT1A line in comparison to its isogenic iPSC-SCPs (Fig. 4a). Using gene ontology (GO) enrichment analysis, we identified pathways regulating the PM component (GO component), receptor signaling and activity (GO function), and cell adhesion (GO process) as the core terms enriched in the CMT1A iPSC-SCPs (Fig. 4b). These data indicate early transcriptional dysregulation in the CMT1A iPSC-SCPs altering the transcriptional networks mediating PM processes, receptor signaling, and cell adhesion, all essential elements for Schwann cell development, differentiation, and maturation.

**Figure 4.**
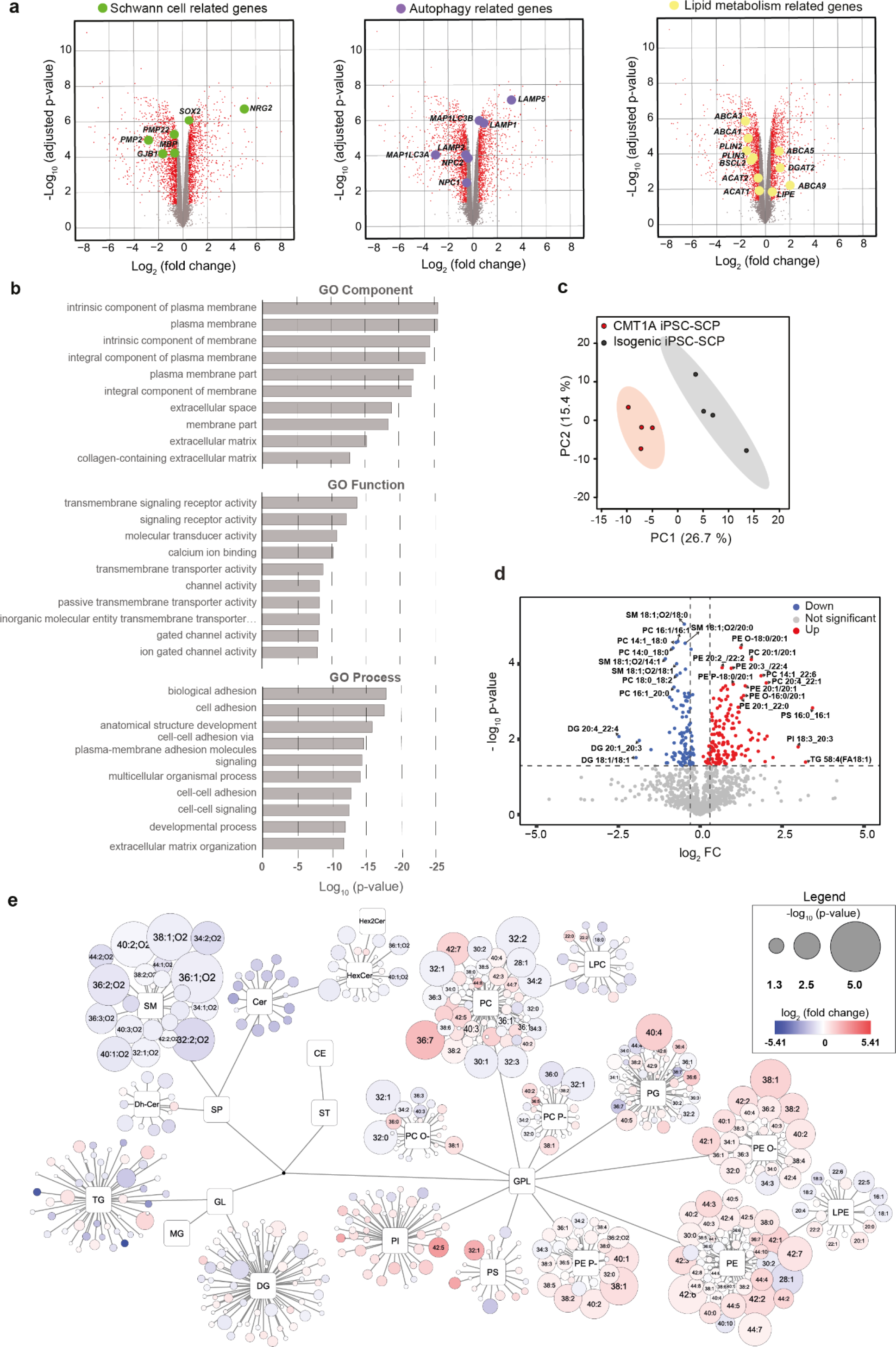
Dysregulated gene expression and alterations in the lipidomic profile in CMT1A patient iPSC-SCPs. **a)** Volcano plots showing dysregulated genes in CMT1A compared to their isogenic control iPSC-SCPs. Dysregulated myelin/Schwann cell-related genes, as well as lipid metabolism and autophagy-related genes, are highlighted in separate graphs. Adjusted P-value threshold was set at 0.05; Fold change (FC) threshold: 0.5. **b)** GO showing the top 10 enriched ontological terms for cellular components (GO Component), function (GO Function), and molecular process (GO Process) that are reduced in CMT1A iPSC-SCPs. **c)** PCA from the lipidomic analysis performed on isogenic and CMT1A iPSC-SCPs. ) Volcano plot of the relative expression profiles of lipid species in the CMT1A iPSC-SCPs. P-value threshold was set at 0.05; log2(FC) threshold: 0.3. **e)** Simplified network visualization of lipid metabolism showing alterations in lipid profiles of CMT1A iPSC-SCPs compared to isogenic controls. Circles represent the detected lipid species, where the circle size expresses the significance according to p-value, while the color darkness defines the degree of up/downregulation (red/blue) according to the fold change. The most discriminating lipids are annotated. Source data are provided as a Source Data file.

**Figure 5.**
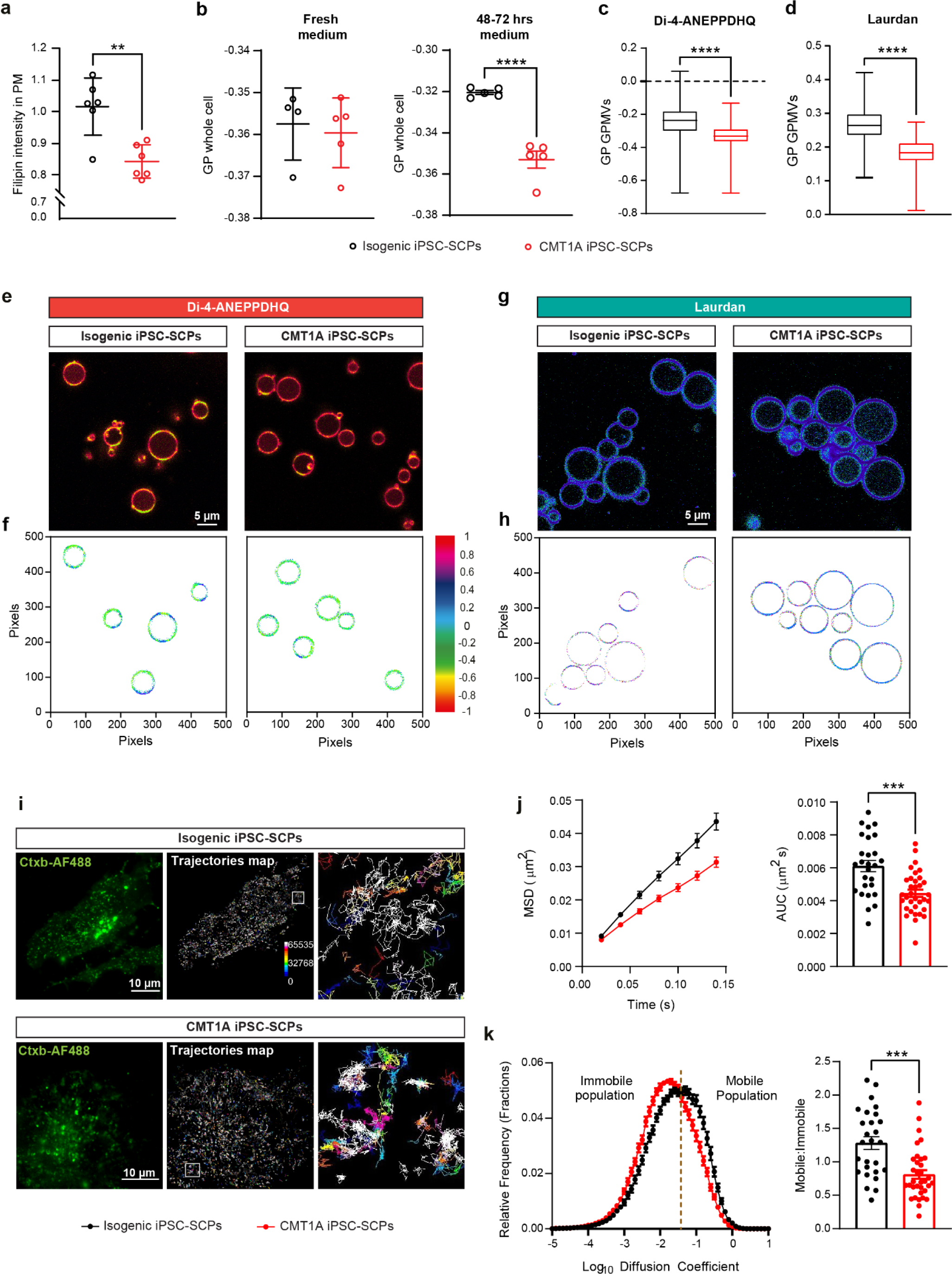
Reduced free cholesterol, increased disorder, and altered lipid raft dynamics in the plasma membrane of CMT1A iPSC-SCPs. **a)** Free cholesterol levels in the plasma membrane of CMT1A iPSC-SCPs and their isogenic controls were determined by a filipin staining. n = average intensity per well, with approx. 200 cells/well analyzed. Two-tailed, unpaired t-test (** p < 0.01). Data are represented as mean ± S.D. **b)** Flow cytometry of Di-4-ANEPPDHQ stained isogenic and CMT1A iPSC-SCPs was used to calculate the GPex value as a measure for membrane ordering [ranging from -1 (low membrane order) to +1 (high membrane order)]. n = individual wells cultured in parallel. Two-tailed, unpaired t-test (**** p < 0.0001). Data are represented as mean ± S.D. **c, d**) GPMVs were generated from isogenic and CMT1A iPSC-SCPs, and used for spectral (lambda, λ) confocal imaging using Di-4-ANEPPDHQ or Laurdan to calculate the GPem in the plasma membrane (see supplemental Fig. 9 + methods). Three independent experiments were performed; representative figures are shown. For Di-4-ANEPPDHQ, CMT1A: n = 164 and isogenic: n = 142 GPMVs, and for Laurdan, CMT1A: n = 151 and isogenic: n = 112 GPMVs analyzed. Two-tailed, unpaired t-test (**** p < 0.0001). **e, f)** Photomicrographs of Di-4-ANEPPDHQ-stained GPMVs from CMT1A and isogenic iPSC-SCPs (e), color-coded GP value per pixel (f). **g, h)** Photomicrographs of Laurdan-stained GPMVs from CMT1A and isogenic iPSC-SCPs (g) color-coded GP value per pixel (h). **i)** Representative TIRF images of cholera toxin b (CTB) labelling in the cell membrane of isogenic and CMT1A iPSC-SCPs, and representative trajectories maps (inset: magnification exhibiting confined and free diffusion). **j-k)** Mean square displacement (MSD) as a function of time, the area under the curve (AUC), the mean distribution of the diffusion coefficient, and the ratio of mobile to immobile fractions for isogenic and CMT1A iPSC-SCPs. Lipid raft dynamics is significantly confined in CMT1A compared to within control isogenic lines. n = 27-34 cells, with 6,966 trajectories per cell in isogenic group and 5,424 trajectories per cells in CMT1A.

**Figure 6.**
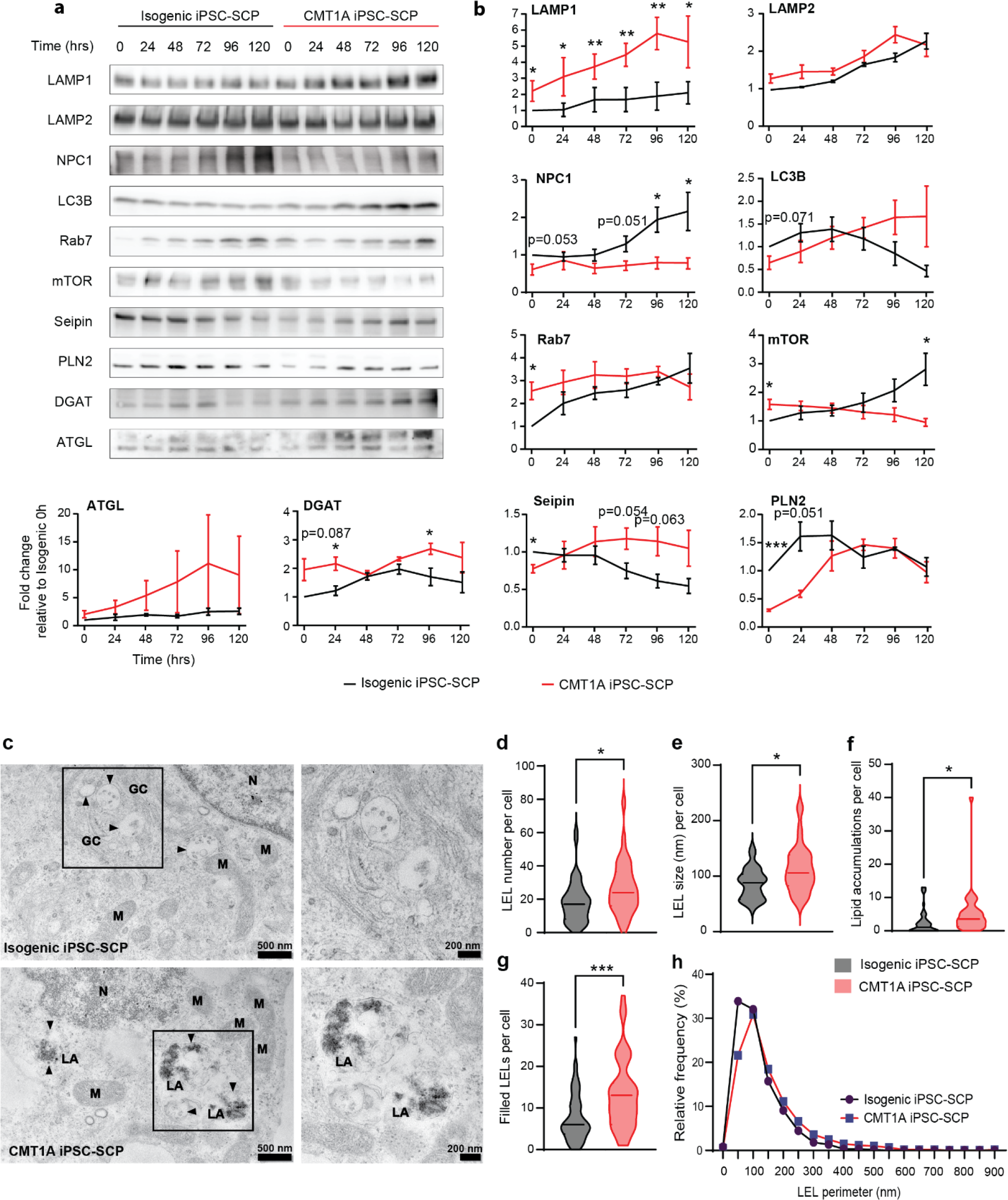
Dysregulated lipid homeostasis and autophagy during lipid stress and lipid accumulations in the late-endosomal lysosomal system in CMT1A iPSC-SCPs. **a)** Immunoblots of key proteins involved in autophagy and lipid homeostasis in isogenic and CMT1A iPSC-SCPs cultured in BSA-free medium (i.e., lipid starvation) over time (0, 24, 48, 72, 96, 120 h). **b)** Quantification of blots displayed in (a). Statistical significance in (b) was evaluated using a two-way ANOVA, followed by a Fisher’s LSD test (* p < 0.05, ** p < 0.01, *** p < 0.001). Data are represented as mean ± S.E.M. n = 3-5 independent time-course experiments. **c)** Transmission electron microscopy (TEM) images of isogenic and CMT1A iPSC-SCPs. M = mitochondria, N = nucleus, GC = Golgi compartment, and LA = lipid accumulation. Black arrowheads indicate the perimeter of the vesicles. A zoomed-in area showing late-endosomal lysosomes (LEL) in isogenic iPSC-SCPs (top), and ‘filled’ LEL with lipid accumulations in CMT1A cells (bottom). **d-g)** TEM quantifications of LEL number (d), and individual LEL size (e) per cell. Lipid accumulations per cell (f). Lipid accumulations were present in LELs and in non-vesicular bound accumulations. The number of ‘filled’ LELs was determined (g). This represents lipid and non-lipid accumulations in LELs that approximately occupied at least 1/5 of the LEL. Isogenic iPSC-SCP cells = 45; CMT1A iPSC-SCP cells = 33. **h)** Relative frequency distribution of the total number of LELs quantified over two independent preparations (isogenic iPSC-SCP LELs = 600; CMT1A iPSC-SCP LEL = 611). Statistical significance in (d-h) was evaluated using a two-tailed, unpaired t-test (* p < 0.05, *** p < 0.001).

**Figure 7.**
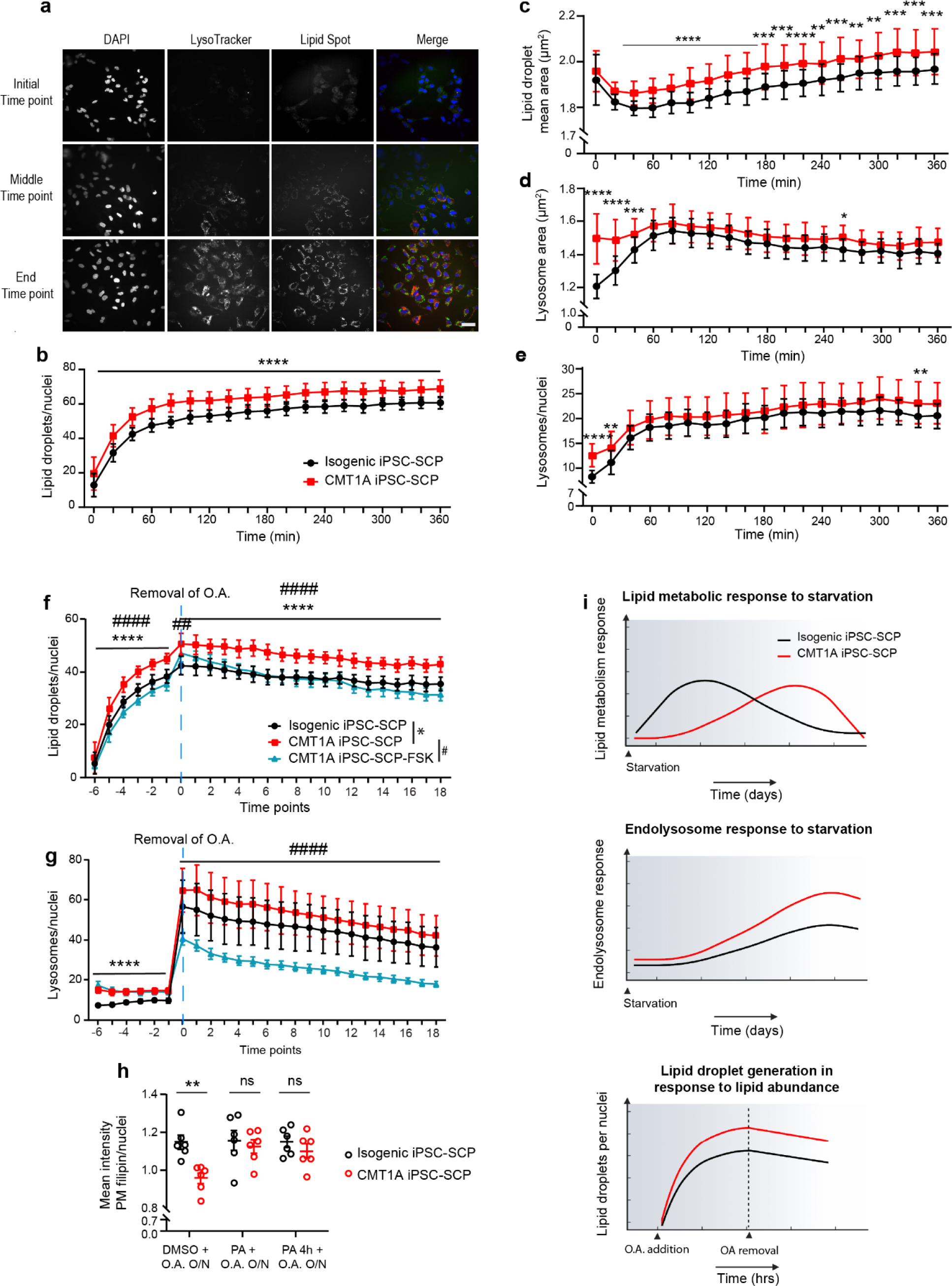
Excessive lipid droplet formation in response to OA treatment in CMT1A iPSC-SCPs which can be reversed by modulating lipid metabolism. **a)** Immunofluorescence images of nuclei (DAPI, in blue), lysosomes (LysoTracker, in red), lipid droplets (LDs, LipidSpot, in green) of CMT1A iPSC-SCPs treated with oleic acid (OA) at the beginning, middle, and end of the time-course experiment. Scale bar = 20 µm. **b-e)** The analysis of the time-course experiment (intervals of 20 min) using an immunofluorescence high-content image analyzer for lysosomes and LDs demonstrated that the CMT1A iPSC-SCPs generate a higher LD number (b) and size (c) in comparison to their isogenic iPSC-SCPs. Also, the lysosome size (d) and number (e) were initially increased in the CMT1A iPSC-SCPs in comparison to their isogenic iPSC-SCPs. Data are represented as LDs or lysosomes/cell/well. 30 wells per group were analyzed for isogenic and CMT1A iPSC-SCPs. Statistical significance was tested by using a one-way ANOVA, with an uncorrected Fisher’s LSD test (** p < 0.01, *** p < 0.001, **** p < 0.0001). Data are represented as mean ± S.D. **f, g)** Time-course experiment using immunofluorescence high-content imaging for lysosomes (lysotracker) and LDs (LipidSpot) demonstrated that the treatment with FSK can prevent the excessive accumulation of LDs (f) and lysosomes (g) in CMT1A iPSC-SCPs during treatment with OA. Time points represent 20 min intervals, and the medium was changed at time point 0, to remove OA from all conditions, but FSK was maintained in the treated condition throughout the experiment. Data are represented as mean ± S.D. **h)** Treatment of CMT1A iPSC-SCPs with the progesterone antagonist (PA) PF-02413873 resulted in increased free cholesterol (filipin intensity) in the plasma membrane. Data are represented as mean ± S.E.M. **i)** An overview of the lipid metabolic and autophagic response of the CMT1A iPSC-SCPs to lipid stress. In f, g, h, data are represented as the number of LD or lysosomes/cell/well. 30 wells per group were analyzed for both isogenic and CMT1A iPSC-SCPs. Statistics in f, g) were performed by using a two-way ANOVA with Šidák multiple comparisons tests ****,^####^ p < 0.0001), and in h by using a two-tailed t-test.

To understand how these alterations in the transcriptome may translate to disturbed lipid profiles of CMT1A iPSC-SCPs, we performed a detailed lipidomic analysis on both the isogenic and CMT1A iPSC-SCP lines. Compared to C3 sciatic nerves, a lower number of lipid species were dysregulated in the CMT1A iPSC-SCPs (Fig. 4d, e). In the PCA score plot, constructed based on the lipid concentrations measured in both cell types, there was a clear separation between CMT1A and isogenic iPSC-SCPs (Fig. 4c). Significantly lower levels of SM were found in CMT1A compared to the isogenic iPSC-SCPs. Additionally, CMT1A iPSC-SCPs had lower levels of Cer, HexCer, and Dh-Cer but, in most cases, without reaching statistical significance. In the CMT1A iPSC-SCPs, many short-chain PC, PC P-, PC O-, PG, PE P-, and PI were downregulated, in contrast, all short-chain PS were upregulated (Fig. 4d, e). We also detected a high accumulation of long-chain (38, 40, 42, and 44 total carbon number) and mostly unsaturated PE O-, PE, and PE P-. Similarly, multiple long-chain and mostly unsaturated PC, PI, and PG species were more abundant in CMT1A (Fig. 4d, e).

#### CMT1A iPSC-SCPs have disordered plasma membranes with reduced lipid raft mobility

Because an altered lipid composition and accumulation of polyunsaturated fatty acids (PUFAs) may affect the fluidity of biological membranes and lipid raft organization and dynamics, we next investigated the membrane properties in the iPSC-SCPs. We analyzed PM levels of free cholesterol of the iPSC-SCPs by performing filipin staining. We detected a significant decrease in the levels of PM cholesterol in CMT1A iPSC-SCPs in comparison to their isogenic control iPSC-SCPs (Fig. 5a). As cholesterol in the PM preferentially interacts with lipids in liquid-ordered (L_o_) domains, such as those that compose lipid rafts, its reduction in CMT1A iPSC-SCPs may affect membrane polarity. To investigate this, cells were labelled with the environmentally sensitive styryl dye Di-4-ANEPPDHQ and analyzed via flow cytometry. The emission spectrum of Di-4-ANEPPDHQ is mainly determined by the cholesterol content in membranes and emits light at 488 nm wavelength in cholesterol-rich L_o_ domains and at 555 nm wavelength in L_d_ domains (Jin et al., 2005). Interestingly, CMT1A iPSC-SCPs membranes became more L_d_ when cultured in the same medium for 48-72h, but not in freshly changed medium (Fig. 5b). This indicates that long-chain fatty acids and PUFAs accumulate in membranes of CMT1A iPSC-SCPs over time.

To specifically explore the PM, we generated giant plasma membrane vesicles (GPMVs) from isogenic and CMT1A iPSC-SCPs. Spectral microscopic analysis on Di-4-ANEPPDHQ labelled GPMVs showed that the generalized polarization (GP) value was significantly lower in CMT1A cells compared to the isogenic control (Fig. 5c, e, f), indicating more L_d_ phases in the PM of CMT1A cells which confirms the flow cytometry data (Fig. 5b). Laurdan, another probe used to study lipid packing that is influenced by the presence of different lipid species (e.g., SM and PC) (Aron et al., 2017; Browning et al., 2020; Sezgin et al., 2015; Suhaj et al., 2020), showed similar reduced GP values in the membrane of CMT1A iPSC-SCPs (Fig. 5d, g, h). Representative images of Di-4-ANEPPDHQ and Laurdan stained GPMVs are displayed, together with the GP value per pixel (Fig. 5e-h). GP histograms clearly indicate an increase in the number of pixels with higher GP values in isogenic GPMVs compared to the CMT1A group (Supplemental Fig. 9c, g). In addition, phasor plots, a model-free Fourier transform analysis approach (Supplemental Fig. 9d, h) (Malacrida et al., 2016; Malacrida et al., 2015; Socas and Ambroggio, 2022; Stefl et al., 2011), demonstrate a significant shift between isogenic and CMT1A GPMVs.

Next, we studied whether the fluidity of the PM was affected in patient-derived SCPs. To do this, lipid raft dynamics in the PM was studied by using live cell total internal reflection (TIRF) imaging with cholera toxin b (CTB, a raft marker) Alexa Fluor 488 in isogenic and CMT1A iPSC-SCPs (Fig. 5 i-k). Single particle tracking was performed, and we found that lipid raft mobility was significantly reduced in the PM of CMT1A iPSC-SCPs. Moreover, the ratio of mobile to immobile raft fractions indicates significantly confined dynamics of raft nanodomains in CMT1A (Fig. 5j, k). Representative TIRF images are shown of CTB AF488 labelling in the cell membrane of isogenic and CMT1A iPSC-SCPs, as well as the trajectory maps illustrating the free hop kind of diffusion in isogenic cells and more confined movement in CMT1A SCPs (Fig. 5i). These data indicate that alterations in CMT1A iPSC-SCP lipidome disrupt the PM properties and lipid raft dynamics.

### Dysregulated lipid homeostasis and autophagy during lipid stress in CMT1A iPSC-SCPs

From both the CMT1A *in vivo* and *in vitro* lipidomic data sets, it is clear that lipid flux is altered. To understand this better, we investigated how prolonged lipid starvation affects lipid homeostasis by monitoring autophagy, lipid biogenesis, and lipolysis over time in CMT1A versus their isogenic control iPSC-SCPs. We deprived the iPSC-SCPs of lipids and collected samples over a time course of 0, 24, 48, 72, 96, and 120 h. We analyzed protein content in iPSC-SCPs and probed the following lysosomal/autophagy markers: lysosomal-associated membrane protein-1 (LAMP1), LAMP2, and microtubule-associated proteins 1A/1B light chain 3B (LC3B). Furthermore, we analyzed the mammalian target of rapamycin (mTOR), a master regulator of autophagy and cellular response to stress and growth, and the late endosome-lysosome (LEL) markers Rab7 and Niemann–Pick C1 protein (NPC1), which are involved in intracellular cholesterol handling and trafficking (Blom et al., 2003; Subramanian and Balch, 2008; Walkley and Suzuki, 2004) (Fig. 6a, b). In the CMT1A iPSC-SCPs, LAMP1 was consistently elevated, while LAMP2 was similarly expressed compared to the isogenic iPSC-SCP levels (Fig. 6a, b). LC3B was progressively upregulated in the CMT1A line, while in the isogenic line, it was initially upregulated followed by a progressive downregulation (Fig. 6a, b). mTOR was initially upregulated in CMT1A iPSC-SCPs, similar to what has been reported in the literature (Krauter, 2021), but throughout lipid starvation, it was downregulated in comparison to isogenic iPSC-SCPs that showed an increase. Rab7 was initially upregulated in CMT1A iPSC-SCPs but failed to mount the same response as the isogenic iPSC-SCPs over time. Interestingly, NPC1 was downregulated throughout the time-course experiment, which suggests alterations in lipid and cholesterol handling and trafficking in LELs in the CMT1A iPSC-SCPs (Fig. 6a, b).

We also investigated markers for LD biogenesis and lipolysis. To do this, we monitored seipin, which is essential for LD biogenesis (Cartwright et al., 2015; Wang et al., 2016), DGAT1, which mediates the conversion of DG and fatty acid-CoA to TG, perilipin 2 (PLN2), and ATGL. PLN2 is correlated to the size of LDs, as it is specifically localized on the LD membrane (Blanchette-Mackie et al., 1995). During lipid starvation, LD biogenesis and size were delayed in the CMT1A iPSC-SCPs, while DGAT1 was slightly elevated at the start and at the end of the time course. ATGL showed a trend of progressive upregulation in the CMT1A iPSC-SCPs. Taken together, these data indicate that CMT1A iPSC-SCPs excessively stimulate the autophagic response and have a delayed LD-mediated response to lipid starvation.

### Lipids accumulate in the late endosome lysosome (LEL) system in CMT1A iPSC-SCPs

In addition to the altered lipid and autophagic flux response, the elevation of the late endosome marker Rab7 and the downregulation of NPC1 at basal conditions in CMT1A iPSC-SCPs could indicate a dysregulated intracellular lipid and cholesterol handling and trafficking. Moreover, this may result in the accumulation of lipid products in LELs. To investigate this, we performed transmission electron microscopy on CMT1A and isogenic iPSC-SCPs. Observable lipid accumulations were prominently visible in LELs of CMT1A iPSC-SCPs compared to controls (Fig. 6c, f, g). Furthermore, LEL number and size were increased in the CMT1A group (Fig. 6d, e, h).

### Increased lipid droplet generation in CMT1A iPSC-SCPs during OA stimulation

As lipid storage and homeostasis are dysregulated in CMT1A iPSC-SCPs during lipid deprivation and prolonged medium incubations, we aimed to understand how CMT1A iPSC-SCPs mobilize lipids during short-term lipid abundance. Therefore, we investigated how the CMT1A iPSC-SCPs respond to the induction of LD formation. Cells were treated with OA and were recorded live over a 7 h period using LipidSpot dye and LysoTracker to monitor the generation of LD and lysosomes, respectively (Fig. 7a). Interestingly, the number and size of LDs were significantly increased in the CMT1A iPSC-SCPs in comparison to their isogenic control (Fig. 7b, c). Additionally, lysosomal size and number were initially upregulated at time-point 0 h (Fig. 7d, e), but then normalized in comparison to the isogenic iPSC-SCPs during continuous OA treatment. These data validate the previous finding that LEL size and number were increased in the CMT1A iPSC-SCPs. Moreover, the excessive response of the CMT1A iPSC-SCPs to OA in the size and production of LD indicates that additional copies of *PMP22* cause abnormal lipid storage and lipid-mediated stress.

### Stimulation of autophagy and lipolysis restored perturbed lipid droplet phenotype and free cholesterol incorporation in CMT1A iPSC-SCP PMs during lipid stress

In order to validate these lipid metabolic phenotypes, we used the CMT1A iPSC-SCPs to investigate whether modulating lipolysis and autophagy improved the LD phenotype present during OA treatment. Therefore, we stimulated CMT1A iPSC-SCPs with forskolin (FSK), a cAMP activator, which is a potent stimulator of lipolysis and autophagy (Duncan et al., 2007; Grisan et al., 2021; Ugland et al., 2011). We observed that FSK reduced LD generation in the presence of OA (Fig. 7f). In addition, once OA was removed, the slope of decline in the reduction of LDs appeared similar between the CMT1A and isogenic iPSC-SCPs. However, FSK continued to stimulate LD breakdown (Fig. 7f). FSK did not have a strong effect on lysosome number in the presence of OA but, once OA was removed, the number of lysosomes was significantly reduced in the FSK-treated CMT1A iPSC-SCPs (Fig. 7g). An overview of the lipid metabolic and autophagic response of the CMT1A iPSC-SCPs to lipid-induced stress is shown in Fig. 7i.

Next, we stimulated free cholesterol incorporation into the PM of the CMT1A iPSC-SCPs by using a progesterone receptor antagonist (PA). Progesterone itself is synthesized from cholesterol in a two-step process (Christenson and Devoto, 2003) and acts as an inhibitor of cholesterol biosynthesis (Metherall et al., 1996). Moreover, progesterone is also a potent activator of PMP22 expression, and the treatment of the CMT1A rat model with a PA was shown to improve the CMT phenotype (Sereda et al., 2003). Treatment with a PA for 4 h or overnight was sufficient to rescue the PM cholesterol deficit in the CMT1A iPSC-SCPs (Fig. 7h).

## Discussion

CMT1A pathophysiology is characterized by a slowly progressive dysmyelinating phenotype starting from early development that leads to axonal loss, denervation of neuromuscular junctions and sensory problems. We used a combination of CMT1A mouse models and patient-derived iPSC-SCPs to investigate how pathological alterations in lipid metabolism cause CMT1A. We identified cholesterol biosynthesis and lipid metabolism as the main canonical pathway downregulated in the sciatic nerves of both the C3 and C22 mouse models. *PMP22* copy number had a dose-dependent effect on cholesterol biosynthesis, which we found by comparing the C3 and C22 mouse models that express 5 and 7 copies of the human *PMP22* gene, respectively. High cholesterol levels are essential for myelin membrane growth (Saher and Stumpf, 2015). Similarly, *PMP22* overexpression dose-dependently reduced the expression of genes regulating the lipidome transcriptional network. In the outbred rat model of CMT1A (Sereda et al., 1996), rats can have varying levels of rodent *Pmp22* expression, giving rise to a spectrum of phenotype severities, including variations in the expression of lipid metabolic-related genes (Fledrich et al., 2018; Fledrich et al., 2012). To understand how these transcriptional changes influence the CMT1A lipid profiles, we performed lipidomics on the sciatic nerves of 5-week-old C3 mice. Overall, the analysis revealed a major downregulation of lipid species in general, but particularly of SPs, with SMs being the most affected in the sciatic nerves of C3 mice compared to their age-matched littermate controls.

The correct PM lipid composition is essential for Schwann cells to interact with axons and to produce myelin, the multilamellar lipid-based structure. During Schwann cell lineage development, neural crest cells give rise to SCPs, that use extending peripheral axons to guide them to the most distal regions of the body (Le Douarin et al., 1991). Once interacting with axons, SCPs can differentiate into immature Schwann cells, which organize axons to be myelinated and non-myelinated in a process defined as radial sorting (Mirsky et al., 2002). To understand how these CMT1A lipid alterations may originate at a cellular level, we developed a protocol to derive a Schwann cell lineage from iPSCs, which was a modified version of Kim et al. (Kim et al., 2017). These differentiated iPSC-SCs had Schwann cell bi- and tripolar morphology, expressed Schwann cell genes, and could myelinate *in vitro*. However, myelination was only seen at an extremely low frequency and could not reliably be used for myelination assays in co-cultures with rodent neurons. Additionally, as there were differences between the CMT1A and its isogenic pair at the final stages of the differentiation, it was difficult to consistently conduct omics analysis on the patient-derived iPSC-SCs. As differentiation defects in the development of mature Schwann cells have been described in CMT1A rodents (Fledrich et al., 2014) and iPSCs (Shi et al., 2018), we hypothesized that early cellular pathologies originate in the CMT1A iPSC-SCPs, as these are the first cell type to express *PMP22* (Mirsky *et al*., 2002). In each iPSC line, *PMP22* copy levels were confirmed using digital droplet PCR. CMT1A iPSC-SCP *PMP22* mRNA was observed to be highly influenced by the lipid concentrations in the medium. *PMP22* mRNA was significantly upregulated in the CMT1A iPSC-SCPs in comparison to the isogenic iPSC-SCPs in high BSA medium but reduced when cultured in low BSA medium. High concentration of BSA occasionally caused cellular clumping and detachment. Due to this and the additional effect high serum levels can have on impairing stem cell differentiation, cells were maintained in low serum concentrations (Valmassoi and Schwarz, 1988; Vincent and Odorico, 2009). From *in vitro* studies, it is known that myelin genes, such as *Mbp* (Byravan and Campagnoni, 1994) and *Pmp22* (Bosse et al., 1999), are regulated by serum levels in the cell culture medium. Interestingly, CMT1A patients can have slightly lower or normal levels of *PMP22* mRNA compared to healthy patients, and *PMP22* mRNA levels do not correlate with disease severity (Hanemann et al., 1994; Katona et al., 2009). In fact, *PMP22* copy number, rather than *PMP22* mRNA, strongly correlates with the CMT1A clinical phenotype (Kim et al., 2015; Martinez Thompson & Klein, 2016).

Transcriptionally, we found a large dysregulation of genes in the CMT1A patient iPSC-SCPs. Several genes relating to Schwann cell myelination and development were downregulated. A large proportion of lipid metabolism genes were similarly downregulated, including the ATP binding cassette subfamily A 1 (ABCA1) and ABCA3, known cholesterol efflux transporters. PMP22 is known to regulate ABCA1-mediated cholesterol efflux (Zhou *et al*., 2019). Knockout of *Pmp22* drastically upregulated ABCA1 and induced LD accumulations in the peripheral nerves of *Pmp22* knockout mice (Zhou *et al*., 2019). In addition, Zhou and colleagues similarly showed that CMT1A patient fibroblasts or Schwann cells overexpressing WT-PMP22 triggered cholesterol sequestration to lysosomes, and reduced ATP-binding cassette transporter-dependent cholesterol efflux (Zhou et al., 2020). In the CMT1A iPSC-SCPs, several genes relating to lipid homeostasis and storage were downregulated. Interestingly, autophagy-associated genes *LAMP1* and *LAMP2* were dysregulated, while *NPC1* and *NPC2* were downregulated. NPC1 and NPC2 are intracellular transporters that act in tandem to remove cholesterol from the lysosomal compartment (Saher and Stumpf, 2015; Subramanian and Balch, 2008; Walkley and Suzuki, 2004), and regulate cholesterol homeostasis through the generation of low-density lipoprotein cholesterol-derived oxysterols (Frolov et al., 2003). Loss of function of NPC1 or NPC2 proteins is known to cause the lysosomal storage disease Niemann-Pick disease, type C1 and 2, respectively (Blom *et al*., 2003).

GO enrichment analysis revealed that genes regulating PM components, transmembrane signaling receptor activity, and biological adhesion were the major GO component, GO function, and GO process enriched in the CMT1A iPSC-SCPs, respectively. These are all essential factors regulating axo-glial interactions (Bhat, 2003; Stassart and Woodhoo, 2021).

Similar to the mice, lipidomics analysis discovered significant alterations in SP, specifically a downregulation in SMs, in the CMT1A iPSC-SCPs compared to their isogenic controls. Interestingly, we also detected an enrichment in Cer and in lipids containing long acyl chain unsaturated fatty acids (e.g., PE O-, PE, PE P-, PC, PI, PG). As the storage and release of cholesterol or PUFAs from LDs can influence the degree of L_o_ and L_d_ domains in the PM (Harayama and Shimizu, 2020), we decided to investigate this. By using a high-content image analyzer, we identified that PM cholesterol levels were reduced in the CMT1A iPSC-SCPs. Moreover, we found that CMT1A iPSC-SCP PMs become more disordered over time. Considering that lipid rafts, which are specialized microdomains of the PM, form through a balanced SM-cholesterol interaction, these data indicate possible impaired lipid raft-mediated signaling and lipid transport, which could result in an increased CMT1A PM disorder and accumulation of PUFAs.

Lipid rafts play a critical role in regulating the activity of membrane receptors and signaling molecules, which were GO terms enriched in the CMT1A iPSC-SCP identified from the bulk RNA sequencing dataset. Lipid raft domains are characterized by a high level of saturation, cholesterol, and SPs, and are relatively resistant to thermal and chemical perturbations [reviewed in (Sezgin et al., 2017)]. Our study showed that there are lower levels of cholesterol and SM in CMT1A mice and iPSC-SCPs, and an increase in PM disorder. These changes would disrupt the normal lipid-protein balance in the lipid raft domains and affect the function of membrane receptors. This may result in altered regulation of membrane receptor signaling and myelination. Our live cell tracking experiments on CTB (raft marker) labelled cells confirmed that the dynamics of lipid raft domains in the PM, and thereby also the fluidity of the PM, is affected in CMT1A iPSC-SCPs.

Of notice, the Cer/SM balance can affect receptor-mediated signal transduction and the integrity of lipid rafts (Taniguchi and Okazaki, 2020). Sphingomyelinase can stimulate the production of Cer from SM which, unlike SM, creates a negative electrical environment for cholesterol, making it easier to remove from the membrane. This means that changes in lipid raft formation can impact cholesterol metabolism, and vice versa (Chang et al, 2006; Yu et al, 2005). The lipidomics results demonstrate a significantly decreased SM in both CMT1A mice and iPSC-SCPs, thus suggesting an impairment of SM metabolism. Interestingly, PMP22 has been shown to preferentially partition to cholesterol-rich L_o_ phase domains (Marinko et al., 2020) and is known to be essential for the formation of lipid rafts (Lee *et al*., 2014), indicating an important regulatory role. Moreover, PMP22 is known to alter the architecture of biological membranes and helps to stabilize them (Mittendorf et al., 2017).

As the lipid composition varies substantially during postnatal development, these data indicate that the overexpression of *PMP22* impacts specific lipid classes, mostly SPs, leading to membrane and lipid storage dysregulation. These changes might play a role in causing CMT1A by altering the initiation of myelination or its structural stability. Interestingly, modulating the lipid and, specifically, the SP pathway in CMT1A has been shown to have positive effects *in vitro* (Visigalli et al., 2020). Moreover, changes in SP metabolism have been discovered to be not just limited to the myelin but can also be found in the biological fluids of CMT1A patients, suggesting a systemic metabolic dysfunction in these individuals. As a result, targeting the SP pathway may have a more widespread effect on the disease. From both CMT1A *in vivo* and *in vitro* lipidomic data sets, lipid flux is altered. To understand how CMT1A iPSC-SCPs mobilize lipids in times of stress, we first investigated how CMT1A iPSC-SCPs respond to lipid starvation throughout 120 h. In alignment with the lipidomic and transcriptomics data, LD growth and biogenesis were delayed, indicating that lipid mobilization was impaired in the CMT1A iPSC-SCPs. mTOR, which is a master metabolic regulator of how cells respond to nutrient abundance and starvation, was initially upregulated in basal conditions, similar to what has been reported recently (Krauter, 2021). Interestingly, mTOR was progressively downregulated over the course of lipid starvation, while it was progressively increased in the isogenic iPSC-SCPs. The LEL marker LAMP1 and the autophagy marker LC3B were excessively stimulated in CMT1A iPSC-SCPs in comparison to their isogenic counterpart. Strikingly, in the CMT1A iPSC-SCPs, NPC1 was downregulated from the start and failed to mount the same response to lipid starvation as the isogenic iPSC-SCPs, indicating possible lipid accumulations in LELs. Using electron microscopy, we indeed found increased LEL size, number, and excessive lipid accumulations in the CMT1A iPSC-SCPs compared to isogenic control iPSC-SCPs.

Alterations in lipid handling and storage may affect the trafficking of lipids to cellular membranes. In yeast, catabolism of LDs regulates rapid membrane expansion (Kurat et al., 2009). To understand how CMT1A iPSC-SCPs handle excess lipids, we used a live cell time-course experiment to study the generation of LDs during OA stimulation. Surprisingly, CMT1A iPSC-SCPs excessively stimulated the size and production of LDs in response to OA treatment. In line with our transmission electron microscopy data, lysosome size, and number were increased at the start of each time-course experiment in CMT1A iPSC-SCPs but then normalized to the isogenic levels upon continuous OA stimulation. Collectively, these data indicate that lipid handling and autophagy are impaired in the CMT1A patient cells.

Finally, to validate these lipid phenotypes as a therapeutic targets, we modulated the phenotype using FSK, a cAMP stimulator which is a known activator of lipolysis (Duncan *et al*., 2007), autophagy (Grisan *et al*., 2021; Ugland *et al*., 2011), and myelin gene expression (Gomis-Coloma et al., 2018; Morgan et al., 1991). We additionally used a PA, which is a known inhibitor of *PMP22* expression (Desarnaud et al., 1998; Sereda *et al*., 2003), and a stimulator of free cholesterol release (Metherall *et al*., 1996). Using the LD-induced phenotype, we demonstrated that FSK could rescue excessive LD production and, when OA was removed from the cell culture medium, FSK treatment significantly reduced the number of lysosomes per cell. Next, as a proof-of-concept, a PA was used to modulate the cholesterol deficit in the CMT1A iPSC-SCPs. As the free cholesterol stain filipin could not be analyzed in live cells due to the necessity to fix them, we analyzed the CMT1A iPSC-SCPs response to 4 h and overnight treatment. After both 4 h and overnight treatment, we found restoration of PM-free cholesterol levels in the CMT1A iPSC-SCPs.

In conclusion, we demonstrated that perturbed cellular lipid homeostasis in CMT1A originates early in the Schwann cell lineage. Moreover, the lipid levels can influence PMP22’s regulation of lipid storage and thus, the availability of lipids and cholesterol to the PM. This dysregulation of membrane lipids causes an increase in PM PUFA incorporation, disordering the PM and impairing lipid raft dynamics. Furthermore, by modulating the lipid composition in the cell culture medium, we identified how membrane lipids critically rely on lipid storage homeostasis, which is strongly altered in CMT1A iPSC-SCPs. These findings highlight PMP22’s role in regulating the plasma membrane’s lipid composition and lipid storage homeostasis, further demonstrating that targeting lipid metabolic pathways is a potential therapeutic intervention for CMT1A patients.

## Experimental procedures

### Animals

Animal experiments were either conducted at Leiden University Medical Center (LUMC) or at KU Leuven. For LUMC: all experiments were done after ethical approval by the institutional ethical committee and the central (national) commission for animal experiments according to EU directive 2010/ 63/EU. The C3-PMP22 mouse model (C3 mice) is a spontaneous revertant of C22 mice (Verhamme *et al*., 2011). C22 and C3 mice were bred heterozygous using WT males x heterozygous females and kept Specific Pathogen Free (SPF) at Janvier-labs (Le Genest Saint Isle, France). Animals were sent to the LUMC or the Free University (Sylics) for experiments and allowed to acclimatize for at least one week. All animals were housed socially with at least 3 mice per cage. The animal room was maintained at a temperature of 21±2 °C, with a relative humidity between 45% and 65%, and a 12-hour light-dark cycle. All cages were provided with sawdust as bedding material, and food (commercial rodent diet) and drinking water were provided ad libitum. For KU Leuven: all procedures were conducted in accordance with the ethical standards for experiments on animals established and approved by the Animal Ethics Committee of KU Leuven. Experiments were conducted under the ethical committee approval n° 104/2017. Breeding was done at KU Leuven and the C3 mice were maintained in a C57BL6/J genetic background. Animals were kept and bred in the same conditions as described above. To verify the genotype and copy numbers of human *PMP22*, digital droplet PCR (ddPCR) was performed as previously described (Prior *et al*., 2022).

### Droplet digital PCR

A qPCR assay probe (6-FAM/ZEN/IBFQ) was designed using Integrated DNA Technology (IDT) RealTime PCR primer design tool based on the location in which the *PMP22* duplication in the isogenic iPSC line was corrected, as documented by (Mukherjee-Clavin et al., 2019). The following human *PMP22* primers and probes were used:

Forward: 5’ – AGG CAG AAA CTC CGC TG – 3’

Reverse: 5’ – ACG AAC AGC AGC ACC AG – 3’

Probe: 5’ – CGA TGA TAC TCA GCA ACA GGA GGC A – 3’

The reference gene used in combination with the designed human *PMP22* assay was the commercially established *AP3B1* (dHsaCP1000001; #10031245, Bio-Rad, Basel, Switzerland) ddPCR copy number variation assay.

### Sciatic nerve immunohistochemistry analysis

Sciatic nerves were dissected from WT, C3, and C22 mice and fixed in 10% formalin for a week. Afterward, tissue was embedded in paraffin, and cross-sections of 5 µm thickness made. Tissue sections were first deparaffinized and rehydrated (from xylene to ethanol 50%, and finally in MilliQ water), and then an antigen retrieval step was performed in a citrate buffer, pH 6.0 (30 min at 96°C). Sections were allowed to cool down in phosphate-buffered saline (PBS) and next the endogenous peroxidase activity was blocked by incubating the sections with 0.3% H_2_O_2_ in methanol for 20 min at room temperature (RT). Afterward, the sciatic nerve sections were blocked for mouse immunoglobulins by using the Dako mouse blocking serum (normal mouse serum in Tris-HCl buffer containing stabilizing protein and 0.015 mol/L sodium azide) for 20 min at RT. Sections were then incubated with the monoclonal mouse anti-human PMP22 antibody (1:100, Abcam, Cambridge, UK, #ab90782) overnight at 4^°^C in a humidified chamber. The following day, sections were washed in PBS and incubated with a goat anti-mouse HRP IgG (1:100, Dako) secondary antibody for 30 min at RT. After several washing steps with PBS, detection of peroxidase activity was achieved with DAB plus (approx. 2-3 min). A Haematoxylin counterstaining was performed to visualize the nuclei. After some final washing steps, the slides were air dried and mounted using Vectashield antifade mounting medium (Vector Laboratories; #H-1000-10).

### iPSC cultures

The CMT1A line, CS67iCMT-n1, and its isogenic control iPSC line, isogenic-CS67iCMT, were generated from fibroblasts isolated from a male CMT1A patient, as described by (Mukherjee-Clavin *et al*., 2019). These cell lines were produced by Cedars-Sinai hiPSC core facility and are available at https://www.cedars-sinai.edu/Research/Research-Cores/Induced-Pluripotent-Stem-Cell-Core-/Stem-Cell-Lines.aspx. Confirmation of *PMP22* duplication was carried out using ddPCR as described above. iPSCs were maintained in Essential 8™ (E8) flex medium (Gibco) with penicillin-streptomycin (1000 U/ml) and cultured as described (Guo et al., 2017; Stoklund Dittlau et al., 2021). In brief, each week colonies were passaged with 0.5 mM EDTA (Invitrogen) diluted in Dulbecco’s phosphate-buffered saline (DPBS; Sigma-Aldrich) and plated on Matrigel LDEV-Free, Growth Factor Reduced (GFR) Basement Membrane Matrix (Corning®). The absence of mycoplasma contamination was routinely checked by PCR.

### Differentiation of Schwann cell precursors from human-iPSCs

The protocol to generate iPSC-SCPs was adapted from Kim et al. (Kim et al., 2017). Briefly, iPSCs were split into small colonies or single cell state and seeded on Matrigel-GFR in E8 flex medium 24 h before initiation of the protocol. RevitaCell supplement (Thermo Fisher Scientific; #A2644501) was added at 10 µl/ml when seeding the cells. iPSC medium was then switched to neural differentiation medium, which was composed of a 1:1 mix of advanced DMEM/F12, GlutaMAX™ Supplement (Thermo Fisher Scientific, #10565018), and Neurobasal™ medium (Thermo Fisher Scientific, #21103049), containing 0.005% BSA (Sigma-Aldrich, #A7979), B27 without vitamin A (1×) (Thermo Fisher Scientific, #12587010), N2 (1×) (Life technology), 0.11 mM β-mercapto-ethanol (Thermo Fisher Scientific), 2 mM Glutamax (Thermo Fisher Scientific), 3 mM CHIR-99021 (Tocris Biosciences), and 20 µM SB 431542 (Tocris Biosciences). After 6 days, the neural differentiation medium was supplemented with 25 ng/mL NRG1 type III (Peprotech, #100-03), which is henceforth denoted as iPSC-SCP differentiation medium (SCPDM).

### Differentiation of Schwann cell precursors to Schwann cell-like cells

For induction of iPSC-SC differentiation, the medium, and the cell plate coating were changed. On day 27 of the protocol, cells are seeded on poly-L-ornithine (PLO, 100 μg/ml) and laminin (5 μg/ml) coated plates in SCPDM. The following day, the medium was switched to iPSC-SC induction medium, which is as follows: DMEM-F12 supplemented with 100 U/ml Pen/Strep, 40 µM SB 431542, 6 µM Chir 99021, 200 nM Retinoic acid (Sigma-Aldrich, #R2625), 20 ng/ml PDGF-bb (Peprotech, #100-14B), 5 µM FSK (STEMCELL Technologies, #100-0249) and 200 ng/ml NRG1 type III. This stage of the protocol is serum-free and small amounts of cell death can occur depending on the iPSC line. On day 30, the medium is changed to the following composition: DMEM-F12 supplemented with 100 U/ml Pen/Strep, 0.005% BSA, 20 µM SB 431542, 3 µM Chir 99021, 100 nM Retinoic acid, 10 ng/ml PDGF-bb, 5 µM FSK and 200 ng/ml NRG1 type III. On day 32, the medium was switched and maintained in the final medium composition: DMEM-F12 supplemented with 100 U/ml Pen/Strep, 0.005% BSA, 5 µM FSK, and 200 ng/ NRG1 type III. iPSC-SCs in this study were assessed at day 35, however, iPSC-SCs can be maintained in culture for several weeks. For long-term cultures, iPSC-SC cultures can be magnetic activated cell sorter (MACS) purified based on the cell surface marker NGFR (data not shown) (Ravelo et al., 2018).

### Transmission electron microscopy on iPSC-SCPs

CMT1A and isogenic iPSC-SCPs were cultured for 48-72 h without any medium change until cells reached 70% confluency. iPSC-SCPs were then fixed using 5% glutaraldehyde (1 h, RT) followed by 2.5% glutaraldehyde (overnight, 4 °C) in 0.1 M sodium-cacodylate buffer. Fixed iPSC-SCPs were scraped in the presence of 0.1 M sodium-cacodylate buffer, centrifuged (200 g, 10 min) and cell pellets were resuspended and incubated in 1.5% agarose for 30 min at 4 °C. The embedded cell pellets were post-fixed in 1% osmium tetroxide (2 h, RT in the dark), rinsed with dH_2_O, and dehydrated in a graded ethanol series (30-100%). Samples were in bloc stained overnight with 4% uranyl acetate in the 70% ethanol step at 4 °C. Finally, after propylene oxide treatment (2 changes 15 min each), pellets were infiltrated with and embedded in 100% epoxy resin for two days (60 °C). Ultrathin sections of 50 nm thickness were post-stained with 3% uranyl acetate in water (10 min), and in Reynold’s lead citrate (2 min). Micrographs were taken with a JEOL JEM1400 (JEOL, Japan) at 80 KV.

Quantification of LELs: Ultrathin sections of iPSC-SCPs were analyzed with magnifications of between 5000-12000× using transmission electron microscopy. If LELs were extensively touching or fusing with other LEL or organelles, perimeter size was not estimated. Images of a cell represent one image plane within a cell and are not representative of the total LELs per the whole cell.

### RNA sequencing

*In vivo*: Tissue was lysed using a Mikro-Dismembrator S (B. Braun Biotech International GmbH, Sartorius group) in TRIzol (Thermo Fischer Scientific). After phenol/chloroform extraction and precipitation, RNA pellet was dissolved in 100 μl buffer RA1 (Machery-Nagel, Duren, Germany). RNA was purified using NucleoSpin RNA XS Kit (Machery-Nagel, Duren, Germany). RNA concentration was measured by using Qubit, RNA-BR Kit (Thermo Fischer Scientific) and quality check was done with a BioAnalyzer RNA Nano Chip (Agilent). cDNA synthesis was performed using NuGEN Ovation RNA-Seq System v2 (7102-A01; NuGEN, San Carlos, CA, USA) followed by purification with the Qiagen MinElute Kit. DNA was sheared to 200 to 400-bp fragments. The DNA was end polished and dA tailed, and adaptors with Bioo barcodes were ligated (Life Technologies). The fragments were amplified (eight cycles) and quantified with a QuBit (Thermo Fisher Scientific). Sequencing was done by Genomescan (Leiden, the Netherlands; https://www.genomescan.nl/) using a Novaseq 6000 PE150 (Illumina).

*In vitro*: iPSC-SCPs were seeded at 1×10^6^ per well in 6 well well-plates and incubated for 48 h in their culture medium as described above. RNA was extracted from the cells using TRIzol™ reagent. RNA concentration and purity were determined using the Nanodrop ND-1000 (Nanodrop Technologies) and RNA integrity was assessed using a Bioanalyzer 2100 (Agilent). Per sample, an amount of 1 µg of total RNA was used as input. Using the Illumina TruSeq® Stranded mRNA Sample Prep Kit (protocol version: Part # 1000000040498 v00 October 2017) poly-A containing mRNA molecules were purified from the total RNA input using poly-T oligo-attached magnetic beads. In a reverse transcription reaction using random primers, RNA was converted into first-strand cDNA and subsequently converted into double-stranded cDNA in a second-strand cDNA synthesis reaction using DNA Polymerase I and RNAse H. The cDNA fragments were extended with a single ’A’ base to the 3’ ends of the blunt-ended cDNA fragments after which multiple indexing adapters were ligated introducing different barcodes for each sample. Finally, enrichment PCR was conducted to enrich those DNA fragments that have adapter molecules on both ends and to amplify the amount of DNA in the library. Sequence libraries of each sample were equimolarly pooled and sequenced on Illumina NovaSeq 6000 (S1 flowcell 100 cycles kit v1.5, 100 bp, Single Reads) at the VIB Nucleomics Core (www.nucleomics.be).

Analysis of the RNA sequencing data was done by using the https://unicle.life/ platform analysis.

### Pathway analysis

RNA-Seq files were processed using the open source BIOWDL RNAseq pipeline v3.0.0 developed at the LUMC. This pipeline performs FASTQ preprocessing (including quality control, quality trimming, and adapter clipping), RNA-Seq alignment and read quantification. FastQC (v0.11.7) was used for checking raw read QC. Adapter clipping was performed using Cutadapt (v2.4) with default settings. RNA-Seq reads were aligned using STAR (v2.7.3a) on the GRCm38 reference genome. The gene read quantification was performed using HTSeq-count (v0.11.2) with setting “–stranded yes”. The gene annotation used for quantification was Ensembl version 99. The raw counts were converted to CPM counts using R package EdgeR (v3.28.1) and from this, fold changes were calculated for every comparison. Data were analyzed through the use of IPA (QIAGEN Inc.)

### Quantitative polymerase chain reaction and transcriptomics

RNA was purified as mentioned above via TRIzol™ reagent. SuperScript™ III First-Strand Synthesis System (Invitrogen) was used to convert RNA (500 ng) to cDNA, according to the manufactures instructions. For qPCR, Fast SYBR^TM^ Green Master Mix (Applied Biosystems^TM^) was used, and the run was performed with the StepOneTM Real-Time PCR System (Applied Biosystems^TM^). The genes of interest obtained from the data were normalized to the reference genes GAPDH and β-actin and further analyzed with qbase+ (Biogazelle). An overview of the primers used in this study is listed in the supplemental section (Supplemental Table 1).

### Preparation of samples for lipidomics

For *in vivo* lipidomic analysis, mice were first anesthetized by an i.p. injection with sodium pentobarbital (200 mg/kg: Dolethal). Then, mice were transcardially perfused with PBS, and following this, sciatic nerves were isolated and flash-frozen in ice-cold isopentane. Samples were then stored at - 80°C until processed by Lipometrix. For *in vitro* lipidomic analysis, cells were seeded at 1×10^6^ density per well in 6 well-plates on day 0. On day 1, the medium was changed. On day 3, the cell culture medium was removed, and the cells were gently washed several times with ice-cold PBS on ice. Cells were then collected in a small amount of ice-cold PBS by scraping and centrifuged at 10,000 Relative Centrifugal Force (RCF) at 4°C. PBS was removed, and pellets were stored at -80°C until processed by Lipometrix.

### Lipidomics

Before the extraction of lipids, cells and mouse sciatic nerves were homogenized in water using an Ultrasonic Tissue Homogenizer UP100H (Hielscher Ultrasonics). For the extraction, an amount of sample containing 10 µg DNA was taken and diluted in 700 μl water. Next, 800 μl of 1 N HCl:CH_3_OH 1:8 (v/v), 900 μl CHCl_3_, 200 μg/ml of the antioxidant 2,6-di-tert-butyl-4-methylphenol (BHT; Sigma Aldrich) solution were added. For the quantitation of lipids, the samples were spiked with 3 μl of SPLASH® LIPIDOMIX® Mass Spec Standard (#330707, Avanti Polar Lipids), 3 μl of Ceramides and 3 μl of Hexosylceramides Internal Standards (#5040167 and #5040398, AB SCIEX). After vortexing and centrifugation, the lower organic phases were collected, and organic solvents were evaporated using a Savant Speedvac spd111v (Thermo Fisher Scientific) at room temperature. The remaining lipid pellet was stored at -20°C under argon.

Just before mass spectrometry analysis, the precipitates were reconstituted in 100% ethanol. Lipid species were determined via liquid chromatography electrospray ionization tandem mass spectrometry (LC-ESI/MS/MS), using the Nexera X2 UHPLC system (Shimadzu) coupled with hybrid triple quadrupole/linear ion trap mass spectrometer (6500+ QTRAP system; AB SCIEX). Chromatographic separation was performed on a XBridge amide column (150 mm × 4.6 mm, 3.5 μm; Waters) maintained at 35°C. As mobile phase, a 1 mM ammonium acetate in water-acetonitrile 5:95 (v/v) was used, and mobile phase B 1 mM ammonium acetate in water-acetonitrile 50:50 (v/v). The following gradient was applied: 0-6 min: 0% B → 6% B; 6-10 min: 6% B → 25% B; 10-11 min: 25% B → 98% B; 11-13 min: 98% B → 100% B; 13-19 min: 100% B; 19-24 min: 0% B. The initial flow rate was 0.7 ml/min and it was increased to 1.5 ml/min from 13 min onwards. SM, Cer, HexCer, and Hex2Cer were measured in positive ion mode with a precursor scan of 184.1, and 264.4 for sphingosine-based Cer, HexCer, and Hex2Cer. TG and DG were measured in positive ion mode with a neutral loss scan for one of the fatty acyl moieties. PC, LPC, PE, LPE, PG, PI, and PS were measured in negative ion mode by fatty acyl fragment ions. Lipid quantification was performed by scheduled multiple reactions monitoring (MRM), the transitions being based on the neutral losses or the typical product ions as described above. The instrument parameters were as follows: Curtain Gas (CUR): of 35 psi; Collision Gas: 8 a.u. (medium); IonSpray Voltage (IS): 5500 V (positive ion mode) and −4500 V (negative ion mode); Temperature (TEM): 550°C; Ion Source Gas 1 (GS1): 50 psi; Ion Source Gas 2 (GS2): 60 psi; Declustering Potential (DP): 60 V and −80 V; Entrance Potential (EP): 10 V and −10 V; Collision Cell Exit Potential (CXP): 15 V and −15 V.

The following fatty acyl moieties were taken into account for the lipidomic analysis: 14:0, 14:1, 16:0, 16:1, 16:2, 18:0, 18:1, 18:2, 18:3, 20:0, 20:1, 20:2, 20:3, 20:4, 20:5, 22:0, 22:1, 22:2, 22:4, 22:5 and 22:6 except for TG which considered: 16:0, 16:1, 18:0, 18:1, 18:2, 18:3, 20:3, 20:4, 20:5, 22:2, 22:3, 22:4, 22:5, 22:6. Peak integration was performed with the MultiQuant^TM^ (v 3.0.3, Sciex). Lipid species signals were corrected for isotopic contributions using an in-house-prepared Python script and library Molmass 2019.1.1. Finally, lipid concentrations were computed, using a one-point standard curve approach. According to the guidelines of the Lipidomics Standards Initiative (LSI), a level 2 type quantification was performed in this way. Two matrices of data were used subsequently in the statistical analysis: quantitative with fatty acyl moieties considered as well as sum notations.

### Miniguide notation for lipid structures

Shorthand notation for lipid structures used in the article followed the proposal built upon LIPID MAPS terminology (Liebisch et al., 2020; Liebisch et al., 2013) (see list of abbreviations).

At the species level, lipid shorthand notations consist of the abbreviation of the lipid class name, followed by the total number of carbons in fatty acyls, colon, and the total number of double bonds. For example, PC 34:1 is phosphatidylcholine with the total number of carbon atoms in both fatty acyls equal to 34 and 1 double bond in the structure. The same rule applies to sphingolipids, with the addition of a semicolon followed by the total number of oxygen atoms (except the amide oxygen). For example, Cer 34:1;O2 refers to a ceramide with a total number of carbon atoms equal to 34, 1 double bond in the structure, and 2 additional oxygen atoms. The molecular species level shows the exact fatty acyl composition; however, sn-1 and sn-2 positions remain unknown, and the “_” separator is used. For example, PC 16:0_18:1, refers to phosphatidylcholine containing FA 16:0 and FA 18:1 attached to the glycerol backbone without knowing their sn-1 or sn-2 positions. In contrast, a “/” separator indicates the exact sn-1 and sn-2 position of constituents (sn-position level). For example, PC 16:0/18:1 refers to phosphatidylcholine with FA 16:0 attached in the sn-1 position and FA 18:1 attached in the sn-2 position, respectively. For sphingolipids, at the molecular species level, the sphingoid base is separated by “/” from the N-linked fatty acid, e.g., Cer 18:1;O2/16:0, indicates a ceramide where a base is a sphingosine containing C18 atoms with one double bond and two hydroxyl groups, and the N-linked fatty acid is FA 16:0.

### Analyzing membrane polarity using flow cytometry and spectral imaging

iPSC-SCPs were seeded at 1×10^6^ per well in 6 well well-plates on day 0. On day 1, the medium was changed. On day 3, iPSC-SCPs were collected for flow cytometry analysis. First, the medium was removed, and cells were stained with Di-4-ANEPPDHQ (Thermo Fisher Scientific; #D36802) at 1:1000 dilution in 1ml PBS. iPSC-SCPs were then incubated for 30 min in a 37°C incubator. iPSC-SCPs were then harvested and washed 3× in PBS, with centrifugation steps of 2 min, 10,000 RCF between each wash step. iPSC-SCPs were resuspended in 500 µl PBS. For flow cytometry, single cells were recorded at excitation wavelengths of fluorescein isothiocyanate (FITC; 495 nm) and phycoerythrin (PE; 566 nm). Values were normalized by subtracting unstained blanks and the excitation generalized polarization (GP_ex_) (Jay and Hamilton, 2017; Parasassi and Gratton, 1995) was calculated as a measure for membrane order, ranging from -1 (L_d_) to +1 (L_o_).

Equation GP_ex_ via flow cytometry

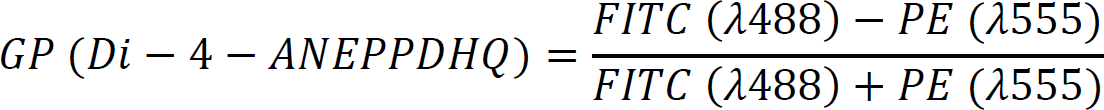

To study membrane polarity via spectral imaging, GPMVs were generated from CMT1A iPSC-SCPs and isogenic controls as described by (Sezgin et al., 2012). Briefly, iPSC-SCPS were seeded at a density of 500,000 cells in T25 flasks and cultured for 72 h as described above. Afterward, the medium was removed, and cells were washed (3×) with PBS and stimulated for 2 h at 37°C with GPMV buffer (10 mM HEPES, 150 mM NaCl and 2 mM CaCl_2_ in MiQ, pH 7.4) supplemented with 25 mM paraformaldehyde (PFA) and 2 mM dithiothreitol (DTT) which are vesiculation forming agents. Next, CMT1A and isogenic GPMVs were collected in the GPMV buffer and labeled by adding either Di-4-ANEPPDHQ (Thermo Fisher Scientific; #D36802) at 1/1000 dilution or Laurdan (#40227, 6-Dodecanoyl-N,N-dimethyl-2-naphthylamine, Merck, Sigma-Aldrich) at 1/500 dilution for 30 min at 37°C in the dark. Following this, the labeled GPMVs were transferred and divided over different wells in an 8-well microslide chamber (Ibidi). GPMVs were allowed to settle to the bottom of the wells for 15 min before imaging was started. On-stage specimen temperature was maintained at 23°C. Spectrally resolved images of the GPMVs were recorded on a Zeiss LSM880 scanhead mounted on the rear port of a Zeiss AxioObserver Z.1 inverted fluorescence microscope equipped with a C-Apochromat 63×/1.2 W Korr M27 water immersion objective lens. Excitation was provided by the 488 nm line of a CW Argon laser used at 0.8% (∼17.5 µW recorded at the stage specimen position) for the Di-4-ANEPPDHQ stained samples. Alternatively, excitation was provided by a tunable MaiTai HPDS DeepSee™ 80 MHz femtosecond pulsed laser set to 810 nm with a requested transmission through the AOM of 3.5% (∼19 mW average power on stage) for the Laurdan stained samples. Care was taken to minimize photobleaching. Excitation and emission light were separated by a MBS488 or a MBS760+ dichroic mirror for the Di-4-ANEPPDHQ and Laurdan samples respectively. Emission was recorded by the 32-channel Quasar spectral GaAsP detector unit over a spectral range of 410-690 nm (8.9 nm spectral channel width). The pixel dwell time was set to 0.85 µs without additional averaging. An area of 45 µm × 45 µm (Zoom 3) was imaged using 512 × 512 pixels resulting in a pixel size of 88 nm × 88 nm and a total acquisition time of 223 ms. The microscope and acquisition configuration were controlled with ZEN Black 2.6 SP1. Based on the collected spectra, corrected for Quasar spectral response (Hoffmann et al., 2020; Velapoldi and Tonnesen, 2004) an emission generalized polarization (GP_em_) value can be calculated as described above. Laurdan and Di-4-ANEPPDHQ are both environmental-sensitive dyes that respond differently depending on the polarity of the membranes by emitting light at different wavelengths (Brewer et al., 2010; Owen et al., 2012; Parisio et al., 2011; Sanchez et al., 2012). A more red-shifted spectrum is correlated with less membrane ordering, while a more blue-shifted spectrum is correlated with higher membrane order (Supplemental Fig. 9). A custom-written Matlab protocol for GPMV data analysis was used to automatically calculate GP values [Based on (Slenders et al., 2018), similar to (Aron *et al*., 2017)]. Briefly, images were processed before corrected spectra were analyzed, which included background subtraction, a Hough transform that selected only fully circular cross-sections displaying proper signal/threshold without “membrane blebbing”, employing openware ImageJ (NIH, Bethesda, MD, USA), and plugins LSM Toolbox, Bio-Formats, as well as academic licensed Matlab (MathWorks®) including Zeiss .czi file reading and the Open Microscopy Environment (OME) script.

Equation GP_em_ via spectral imaging on a confocal system:

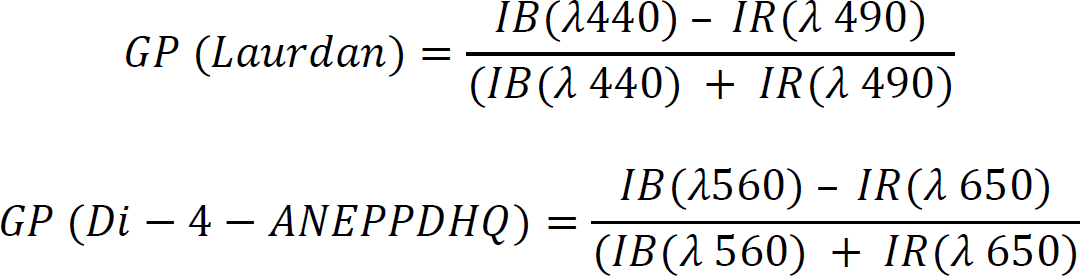

### Lipid raft dynamic assay and analysis

CMT1A and isogenic iPSC-SCPs were cultured in 8 well Ibidi microslide chambers at a density of 5000 cells/well as described above. After 72h in culture in the same medium, lipid rafts were labeled using a Vybrant™ Alexa Fluor 488 lipid raft labeling kit (V34403, Thermo Fisher Scientific) according to the manufacturer’s instructions. After labeling with CTB (which binds ganglioside GM1 in rafts), medium was refreshed, and cells were allowed to recover for 2 hours before live cell imaging was performed. Time series of the dynamics of AF488 labeled CTB in lipid rafts were recorded on a Zeiss Elyra PS.1 inverted widefield fluorescence microscope equipped with an automated TIRF module and an alpha Plan-Apochromat 100×/1.46 DIC M27 oil immersion objective lens. Excitation was provided by a 488 nm CW laser used at 0.5% (∼200 µW recorded from the objective) in objective-based TIRF mode. The TIRF angle was optimized for contrast for each sample (ranging from 65-66°). Excitation and emission light were separated by an MBS 488 dichroic mirror and emission was additionally filtered by a BP495-575 + LP750 emission filter before being recorded by an Andor iXon+ 897 EMCCD camera (512×512 pixels or 16 µm by 16 µm frame size) operating in 16-bit mode with an EM gain of 200 and an exposure time of 33 ms. Additional magnification was provided by a 1.6× optovar lens, resulting in a total magnification of 160× and a pixel size of 100 nm × 100 nm. For each acquisition series, 1000 frames were recorded without delay resulting in a total acquisition time of 33 seconds. The microscope and acquisition configuration were controlled with ZEN Black 2.3 SP1.

Single particle tracking (SPT) on images of the time series of isogenic vs CMT1A iPSC-SCPs was performed by using the Metamorph® software (Molecular Devices) with PALMTracer plugin as described before (Bademosi et al., 2017; Kechkar et al., 2013), in order to quantify and obtain the MSD, diffusion coefficient, and plot highly resolved trajectory maps. In brief, trajectories of localisations that appear in eight or more frames are precisely characterized by PALMTracer. For a trajectory of N data points (coordinates *x(t), y(t)* at times *t* = 0 to (*N* -1) × Δ*t*, (the inverse of the acquisition rate), the MSD for time intervals *τ = n* × Δ*t*) is calculated using the formulae:

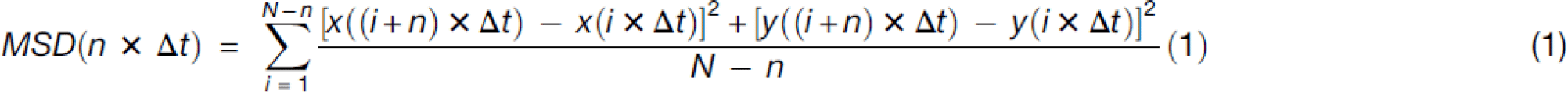

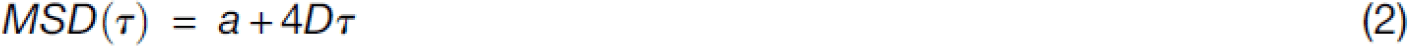

For each MSD, the diffusion coefficient (*D*) is calculated from the linear fits of the first four points by the equation MSD(*t*) = *a* + 4*Dt*, where *a* is the *y*-intercept. The MSD measures the “area” (µm^2^) covered by each molecule over time (33 ms) between frames of the acquired movie. The diffusion coefficient is presented as a relative frequency distribution histogram (Log_10_ of µm^2^ s^−1^).

### Immunocytochemistry

For membrane protein stains, cells were fixed with 4% PFA for 20 min at RT and then washed (3×) in DPBS with calcium and magnesium (DPBS +/+, Gibco Invitrogen). For structural proteins, cells were fixed with ice-cold methanol at -80°C for 10 min, followed again by washing (3×) with DPBS +/+. Cells were then blocked and permeabilized using 5% normal donkey serum (NDS, Sigma-Aldrich), 5% goat serum (Sigma-Aldrich), and 0.1% Triton X-100 (Acros Organics™) in PBS for 30 min. Primary antibodies were diluted 1:50-1:1000 in blocking buffer (2% NDS in PBS) overnight at 4°C (Supplemental Table 2). The following day, after washing (3×) with DPBS +/+, incubation with the appropriate secondary antibodies labeled with either Alexa Fluor 488, 555, or 647 (Invitrogen) at 1:1000 dilution was performed for 2 h at RT. Cells were incubated with NucBlue® (Thermo Fisher Scientific, #R37606) or NucRed® (Thermo Fisher Scientific, #R37106) stain in DPBS for 20 min, washed (3×), and mounted on glass slides using ProLong® Gold antifade reagent without DAPI (Invitrogen, #P36934). Images were acquired using the DMi8 (Leica) confocal microscope. Images were analyzed using the widely used non-commercial software ImageJ FIJI (NIH, Bethesda, MD, USA).

### Western blot analysis

Cells were maintained in the same medium for 48 h prior to collection. For time-point analysis, iPSC-SCPs were seeded at day 0 in SCPDM, and the following morning, the medium was switched to a BSA-free SCPDM. Time-point 0 h was collected at this moment; thereafter, cells were collected every 24 h for a total of 120 h. For protein collection, cells were first washed with ice-cold DPBS and immediately lysed in RIPA buffer (Sigma-Aldrich) supplemented with cOmplete™ Mini, EDTA-free Protease Inhibitor Cocktail (Roche) according to the manufacturer’s instructions. Determination of protein concentration, loading of the Western blot, and subsequent analysis were done as previously described (Benoy et al., 2018; Prior *et al*., 2022). Each marker investigated was normalized to the total protein stain. All blots used for the quantifications are shown in Supplemental Fig. 10 with their corresponding total protein-stained blots that were used for normalization.

### High throughput imaging

The Operetta system (Perkin Elmer) was used for the high throughput image analysis. Cells were seeded on a 96-well plate on day 0 and staining occurred on day 2. For filipin stain, the protocol here reported was followed on the same day of the image acquisition. Cells were washed and incubated for 5 minutes at 37°C with 1X Pre-Staining Solution (MemBrite® Fix 543/560, #30094, Biotium) diluted in DPBS buffer. After a DPBS wash, freshly prepared 1X MemBrite® Fix Dye 543/560 solution (#30094, Biotium) was added for 5 minutes at 37°C. Cells were then washed with DPBS (+/+) and fixed in 4% PFA for 20 min. Cells were then blocked in filtered 5% donkey serum (#D9663, Sigma) for 1 h. Filipin was diluted 1/75 in filtered blocking serum and applied for 2 h at RT in dark. Cells were then washed once in DPBS (+/+) and subsequently incubated with NucRed® for 20 min followed by DPBS (+/+) wash steps (x3). Cells were imaged immediately after staining using the Operetta system and image analysis was performed via the provided Harmony software. Cells were segmented with the help of the digital phase contrast images and PM filipin staining intensity was analyzed. For LD and lysosome accumulation, DAPI, LipidSpot, and LysoTracker, were incubated with cells 30 min prior to imaging. Just before imaging started, the medium was replaced with SCPDM made with DMEM/F12 phenol red-free medium (Thermo Fisher Scientific; #21041025) with oleic acid (OA) (concentration ranged from 10-500 µM for different experiments). Half the normal concentration of LipidSpot and LysoTracker (Supplemental Table 2) were added to the medium during live cell imaging. Analysis of high throughput imaging was done using the Harmony software, through which LDs and lysosomes number and size could be evaluated over time.

### Myelination assay

Mouse DRG neurons were isolated as described (Richner et al., 2017). Once DRGs were removed and placed in ice-cold neuronal growth medium (Neurobasal medium, B27, and 10 ng/ml Nerve growth factor (NGF)-2.5 s (Merck Millipore), 2 mM L-glutamine, and Pen/Strep (50 U/ml)) with 1/100× RevitaCell supplement. Using sterile fine forceps, DRGs were placed gently one at a time on coverslips previously coated with PLO and laminin and completely dried overnight at 37°C in a 5% CO2 incubator (sterile). Each DRG was encapsulated in drops of 50 µl matrigel 3:1 medium mix. Matrigel was allowed to dry for 1 h at 37°C in a 5% CO2 incubator. Next, the neuronal growth medium was added to the DRGs and incubated overnight at 37°C in 5% CO_2_. The medium was then supplemented with cytosine arabinoside (ara-c) to remove contaminating cells and replaced every 24 h for 10 days straight. Wells were checked to ensure no contaminating cells were present before seeding. iPSC-SCs were then seeded on DRGs and maintained in 50:50 neuronal medium and day 35+ iPSC-Schwann cell medium. After 7 days of co-cultures, the medium was switched to a promyelinating medium which contained 5% FBS, with ascorbic acid and FSK removed from the medium. The medium was changed 50% every other day and maintained for at least 2 months.

### Statistics

Most statistical analyses were performed using GraphPad Prism software version 9 (GraphPad Software Inc., California, USA). All data were first checked for normality to select the appropriate statistical test. Unless stated otherwise, the unpaired two-tailed Student *t*-test was used for the comparison of two means and the one-way ANOVA test was performed for the multiple-group analysis.

For GPMV phasor plots, programming in R (https://www.r-project.org/; v 4.2.2) was done to perform MANOVA analysis to obtain data on group distribution (isogenic vs CMT1A) and the Eta squared. For raft dynamics, the Welch-corrected two-tailed *t*-test was used to compare the area under the curves (AUCs) and the mobile:immobile fractions. Statistical analysis for the lipidomics data was performed using the MetaboAnalyst platform (v 5.0) (Pang et al., 2021), R programming language for statistical computing and graphics (v 4.2.2), and Cytoscape software (v 3.9.1). Graphics were merged using Inkscape open-source vector graphics editor (v 1.2). Before performing statistical computations, all matrices were filtered, and lipids with more than 60% missing concentrations were removed. Next, the missing values were substituted for each lipid separately by 80% of the lowest concentration measured. Means with standard deviations were computed for each group using rstatix library (v 0.7.1). The Welch *t*-test was used for comparing means between CMT1A versus isogenic cells and CMT1A versus WT mice. Fold change was computed as a ratio of the mean value in CMT1A to the mean value in isogenic (cells) or WT (mice). Bubble plots reflecting simplified lipid metabolism were generated with the Cytoscape software. In both bubble plots, raw p-values from the Welch *t*-test were presented as -log_10_ and fold change values as log_2_. Volcano plots were prepared using EnhancedVolcano library in R (v 1.16.0) with the cut-off points of 0.3 for the log_2_ fold change (CMT1A to isogenic or CMT1A to WT) and 0.05 raw p-value from the Welch *t*-test (rstatix library). PCA plots were designed using the MetaboAnalyst 5.0 platform after the log_10_-transformation of concentrations and Pareto-scaling.

Data from are presented either as means ± standard error of the mean (S.E.M) or the standard deviation (S.D.). Statistical significance was set at the following: * p < 0.05, ** p < 0.01, *** p < 0.001, and **** p < 0.0001. All figures were modified and finalized in Adobe illustrator (version 2022).

## Acknowledgments/funding

This work was supported by VIB, the KU Leuven (C1 and “Opening the Future” Fund), the “Fund for Scientific Research Flanders” (FWO), the Thierry Latran Foundation, the “Association Belge contre les Maladies neuro-Musculaires” (ABMM), the Muscular Dystrophy Association (MDA), the Association Française les Myopathies (AFM) and the ALS Liga Belgium (“A Cure for ALS”). R. Prior is a Strategic Basic (SB) PhD fellow at Fonds Wetenschappelijk Onderzoek (FWO-Vlaanderen), project number 1S59317N. R. Prior was also supported by the National University of Ireland, Travelling Studentship. A. Silva is supported by the FWO fundamental research PhD program, project number 11N2222N. T. Vangansewinkel is also supported by the FWO fundamental research postdoc program, project number 12Z2620N. FAM is supported by an NHMRC fellowship APP1155794 and idea grant APP2010901. P.V.D. holds a clinical investigatorship of the FWO and is supported by the E. von Behring Chair for Neuromuscular and Neurodegenerative Disorders, the Fund ‘Een Hart voor ALS’ and the ’Laevers Fund for ALS Research’. L.V.D.B. is supported by the Generet Award for Rare Diseases. I. Michailidou is funded by the Prinses Beatrix Spierfonds, grant W.OR14-09. T. Hellings is funded by the Prinses Beatrix Spierfonds, grant W.OB21-02. We thank Lipometrix, the lipidomics core facility of KU Leuven, for the extraction and analysis of the lipidomics datasets and the excellent technical help (https://www.lipometrix.be).

The authors would like to thank Prof. Stein Aerts and Seppe De Winter (KU Leuven-VIB) for their input on bioinformatics analysis. We would like to thank Prof. Catherine Verfaille (KU Leuven) for her initial input on the iPSC-SC protocol. Lastly, we would like to thank Cedars-Senai and Prof. Robert Baloh for the development of the iPSC lines used and for initial input into the story.

## Author contributions

RP conceptualized and designed the study, with input from TV, KF, FB, EW and LVDB. RP developed the iPSC-SC protocol and performed associated experiments with technical help from CM, DVDB, SV and KE. RP collected samples for lipidomics experiments and was processed by JD and JS. RP, AS and KI performed lipidomics analysis. RNA sequencing was conducted with CA, TK, and HM. RP and AS conducted the Western blot and lipid droplet experiments and performed the associated analyses. RP conducted iPSC-SCP bulk RNA sequencing analysis, with input from AKR. AKR performed TEM with assistance from RP and AS. RP performed TEM analysis. VDL performed Di-4-ANEPPDHQ FACS analysis. TV performed all Di-4-ANEPPDHQ or Laurdan GMPV and lipid raft mobility experiments; TV performed the subsequent analysis in coordination with MV, AB and FAM. ER and WV performed initial lysosome experimentation on iPSC-SCPs. RP conducted ddPCR analysis. RP and DVDB performed ICC imaging. FB and KF supervised and coordinating work in relation to the C3 and C22 mice. TPH, IM, and JV performed histological experiments and analysis on work relating to the C3 and C22 mouse models. ML, BK, NV, and NS performed *in vivo* follow-ups of the C3 mice. PVD, JS, IL, FB, KF, EW, and LVDB provided insight and critical feedback throughout the project. RP wrote the manuscript and revised and finalized the manuscript with TV, AS, EW, and LVDB. All authors participated in the finalizing of the manuscript.

## Declaration of interests

The authors declare to not have a conflict of interest.

## Data Availability

All relevant data of the present manuscript are available from the corresponding authors on reasonable request.

## Supplemental section

### List of abbreviations

ABC: ATP-binding cassette transporters
ACAT2: Acetyl-CoA acetyltransferase 2
ACTB: β-actin
Ara-c: Cytosine arabinoside
ATGL: Adipose triglyceride lipase
AUC: Area under the MSD curve
BIII-TUB: βIII-tubulin
BSA: Bovine serum albumin
C3: C3-PMP22
cAMP: Cyclic adenosine monophosphate
CDH19: Cadherin 19
Cer: Ceramide
CMT: Charcot-Marie-Tooth disease
CMT1A: Charcot-Marie-Tooth disease type 1A
CTB: Cholera toxin b
CYP51A1: Cytochrome P450 family 51 subfamily a member 1
ddPCR: Digital droplet PCR
DHH: Desert hedgehog
DRG: Dorsal root ganglion
DG: Diacylglycerol
DGAT1: Diacylglycerol acyltransferase 1
dhCer: Dihydroceramide
DHCR7: 7-dehydrocholesterol reductase
DHCR24: 24-dehydrocholesterol reductase
EBP: EBP cholestenol delta-isomerase
EGR2/KROX-20: Early growth response 2
FDFT1: Farnesyl-diphosphate farnesyltransferase 1
FDPS: Farnesyl pyrophosphate synthase
FSK: Forskolin
GAP43: Growth associated protein 43
GFAP: Glial fibrillary acidic protein
GL: Glycerolipids
GO: Gene ontology
GP: Generalized polarization
GPL: Glycerophospholipids
GPMVs: Giant plasma membrane vesicles
HADHB: Hydroxyacyl-CoA dehydrogenase trifunctional multienzyme complex subunit beta
Hex2Cer: Di-hexosylceramides
HexCer: Mono-hexosylceramides
HMGCR: 3-hydroxy-3-methylglutaryl (HMG) coenzyme A (CoA) reductase
HMGCS: 3-Hydroxy-3-Methylglutaryl-CoA Synthase
HNK-1: Human Natural Killer 1
HSD17B7: Hydroxysteroid 17-beta dehydrogenase 7
IDI1: Isopentenyl-diphosphate delta isomerase 1
iPSC: Induced pluripotent stem cells
iPSC-SCs: Induced pluripotent stem cell-derived Schwann cells
iPSC-SCPs: Induced pluripotent stem cell-derived Schwann cell precursors
ITGA4: Integrin subunit alpha 4
LAMP1: Lysosomal-associated membrane protein-1
LAMP2: Lysosomal-associated membrane protein-2
LBR: Lamin B Receptor
LC3B: Microtubule-associated proteins 1A/1B light chain 3B
LD: Lipid droplet
Ld: Liquid-disordered phase
LEL: Late endosome-lysosome
Lo: Liquid-ordered phase
LPC: Lysophosphatidylcholines
LPE: Lysophosphatidylethanolamines
LSD: Least significant difference
LSS: Lanosterol synthase
MBP: Myelin basic protein
MPZ: Myelin protein zero
MSD: Mean square displacement
mTOR: Mammalian target of rapamycin
MSMO1: Methylsterol monooxygenase 1
MVD: Mevalonate diphosphate decarboxylase
MVK: Mevalonate kinase
NGFR: Nerve growth factor receptor
NPC1: Niemann-Pick C1 protein
NRG1, type III: Neuregulin-1, type III
NSDHL: NAD(P)H steroid dehydrogenase-like protein
OA: Oleic acid
OCT3/4: Octamer-binding transcription factor 3/4
PBS: Phosphate buffer saline
PC: Phosphatidylcholines
PCA: Principal component analysis
PC O-: Phosphatidylcholine ethers
PC P-: Phosphatidylcholine plasmalogens
PDGF?: Platelet-derived growth factor subunit?
PE: Phosphatidylethanolamines
PE O-: Phosphatidylethanolamine ethers
PE P-: Phosphatidylethanolamine plasmalogens
PG: Phosphatidylglycerols
PI: Phosphatidylinositols
PLN2: Perilipin 2
PM: Plasma membrane
PMP22: Peripheral myelin protein 22 (human)
Pmp22: Peripheral myelin protein 22 (mouse)
PMVK: Phosphomevalonate kinase
PNS: Peripheral nervous system
PUFAs: Polyunsaturated fatty acids
RA: Retinoic acid
Rab7: Ras-related protein 7
RT: Room temperature
S100?: S100 calcium-binding protein?
SC5D: Sterol-C5-desaturase
SCP: Schwann cell precursor
SM: Sphingomyelins
SP: Sphingolipids
SQLE: Squalene Epoxidase
SOX2: Sex determining region of Y-related high mobility group box 2
SOX10: Sex determining region of Y-related high mobility group box 10
TFAP2: Transcription factor AP-2?
TG: Triacylglycerol
TIRF: Total internal reflection fluorescence
TM7SF2: Transmembrane 7 Superfamily Member 2
WT: Wild-type<colcnt=1>

## Supplemental Tables

**Table S1:**
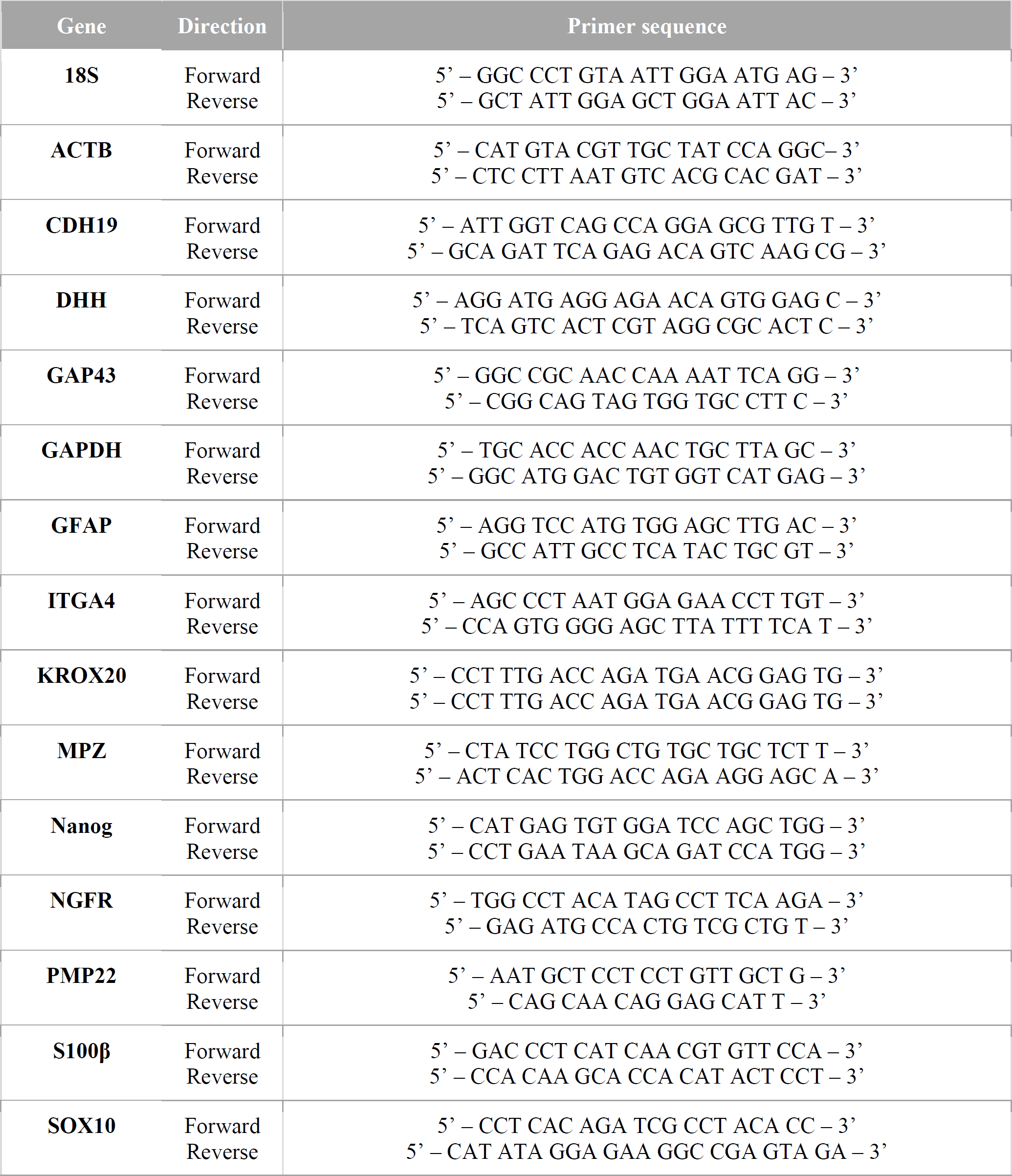
Primers used for qPCR analysis.

**Table S2.** List of primary antibodies and live cell imaging dyes used in this study. FACS = fluorescence activated cell sorting; ICC = immunocytochemistry; IHC = immunohistochemistry; WB = Western blot; SpCI = spectral confocal imaging.

| Antibody | Method | Dilution | Company | Catalogue number |
| --- | --- | --- | --- | --- |
| TFAP2 | ICC | 1/500 | Sigma-Aldrich | HPA028850 |
| ATGL | WB | 1/500 | Santa Cruz | SC365278 |
| BIII-Tubulin | ICC | 1/500 | Abcam | ab7751 |
| Di-4-ANEPPDHQ | FACS,<br>SpCI | 1/1000 | Thermo Fisher<br>Scientific | D36802 |
| DGAT1 | WB | 1/2000 | Abcam | ab181180 |
| EGR2/Krox20 | ICC | 1/300 | NovusBio | NBP2-45696 |
| Filipin | ICC | 1/75 | Sigma-Aldrich | F4767-5MG |
| GAP43 | ICC | 1/100 | Abcam | ab75810 |
| GFAP | ICC | 1/500 | Agilent | z033429-2 |
| HNK-1 | ICC | 1/100 | Sigma-Aldrich | C6680 |
| Laurdan | SpCI | 1/500 | Sigma-Aldrich | #40227 |
| LAMP1 (H4A3) | WB | 1/1000 | DSHB | H4A3 |
| LAMP2 (H4B4) | WB | 1/500 | DSHB | SC-18822 |
| LC3B | WB | 1/2000 | Cell Signaling | 3868S |
| LipidSpot™ 488 | ICC | 1/1000 | Biotium | 70065 |
| LysoTracker™555 | ICC | 1/1000 | Thermo Fisher<br>Scientific | L12492 |
| MBP | ICC | 1/500 | Sigma | ab9348 |
| MemBrite® 543/560 | ICC | 1/1000 | Biotium | 30094-T |
| MPZ | ICC | 1/1000 | NovusBio | NB100-1607 |
| mTOR | WB | 1/500 | Cell Signaling | 2983S |
| Nanog | ICC | 1:300 | Thermo Fisher<br>Scientific | PA1-097X |
| NPC1 | WB | 1/2000 | Abcam | ab134113 |
| OCT4 | ICC | 1/100 | Santa Cruz | Sc-9081 |
| NGFR | ICC | 1/500 | Abcam | ab3125 |
| Perilipin-2/ADFP | WB | 1/500 | NovusBio | NB110-40877 |
| PMP22 | IHC | 1/100 | Abcam | ab15506 |
| PMP22 | ICC | 1/100 | Abcam | ab90782 |
| Rab7 | WB | 1/1000 | Cell Signaling | 9367S |
| S100β | ICC | 1/100 | NovusBio | NBP2-59618 |
| Seipin | WB | 1/500 | Abcam | ab106793 |
| SOX10 | ICC | 1/100 | Abcam | ab155279 |
| SOX2 | ICC | 1/500 | Merck Millipore | ab5603 |

## Supplemental methods

### Semi-thin sectioning

Semi-thin cross-sections were prepared from sciatic nerves of 6-day or 1-year-old C3-PMP22 mice and WT littermates and analyzed as previously described (Prior *et al*., 2022).

### Functional testing

C3-PMP22 mice and their WT littermates were evaluated longitudinally at the following ages: 3, 7, 12, 15, and 19 weeks old. Primary functional outcome measures were tested in these animals at the selected time points: the strength of all front and hind paws combined were measured in the grip strength test, as previously reported (Prior *et al*., 2022). Also, the balance beam test was used to score the ability of mice to traverse a stationary horizontal rod and measure sensorimotor coordination as assessed by the latency to cross the beam and the number of foot slips. Mice were placed at a platform at the start of a wide training beam (100 cm long, 5 cm wide) and allowed to walk along into an enclosed box. All mice had three training runs on a wide beam, and on the next day were given 3 runs on a narrow beam (1 cm wide). A Phenotyper automated home cage (Noldus IT, Wageningen, Netherlands) was used to monitor the spontaneous behavior and free movement of the animals during the night (active phase). Mice were housed for 3 days in the Phenotyper automated home-cage, with access to water, food, and shelter, without intervention. The long movement threshold and the maximum velocity of these long movements were measured. Key kinematic parameters: mice make short movements, such as turning or rearing against the wall, as well as long movements when mice travel from one location in the cage to the next. These two types of movement, i.e., short, and long were distinguished by plotting the frequency distribution of all move segment distances over three days in the cage, yielding a bimodal distribution. An animal-centered threshold was defined by the intersection of the two Gaussian distance curves obtained after the Gaussian mixture model fitting of data of each individual mouse. An animal-centered threshold was set distinguishing the 90% of shortest arrests (short) from the 10% of longest arrests (long).

### Comparing bulk RNA-sequencing data and single-cell atlas of human developing spinal cord

Bulk RNA-sequencing reads were mapped against the hg38 genome using STAR (v2.7.3a) with default parameters. From the mapped reads a count matrix over genes was generated using featureCounts from the Subread (v1.6.3) package with default parameters. The count matrix of the single-cell atlas of developing human spinal cord along with cell metadata containing cell type annotations (Rayon et al., 2021) was downloaded from geo using accession number GSE171890. The single-cell atlas was filtered for annotated cells. To assess transcriptional similarities between the cell types from the single-cell atlas and the bulk RNA-sequencing samples a regression model was fitted with the pseudobulked expression profile from the single-cell atlas as the dependent variable and the bulk RNA-sequencing expression profiles as independent variables, for each cell type in the single cell atlas, using non-negative least squares regression (Scikit-learn, v0.24.1). The regression coefficients from these models were interpreted as a measurement of similarity between the bulk RNA-sequencing samples and the transcriptional profiles of the cell types from the single-cell atlas.

## Supplemental Figures

**Supplemental Figure 1.**
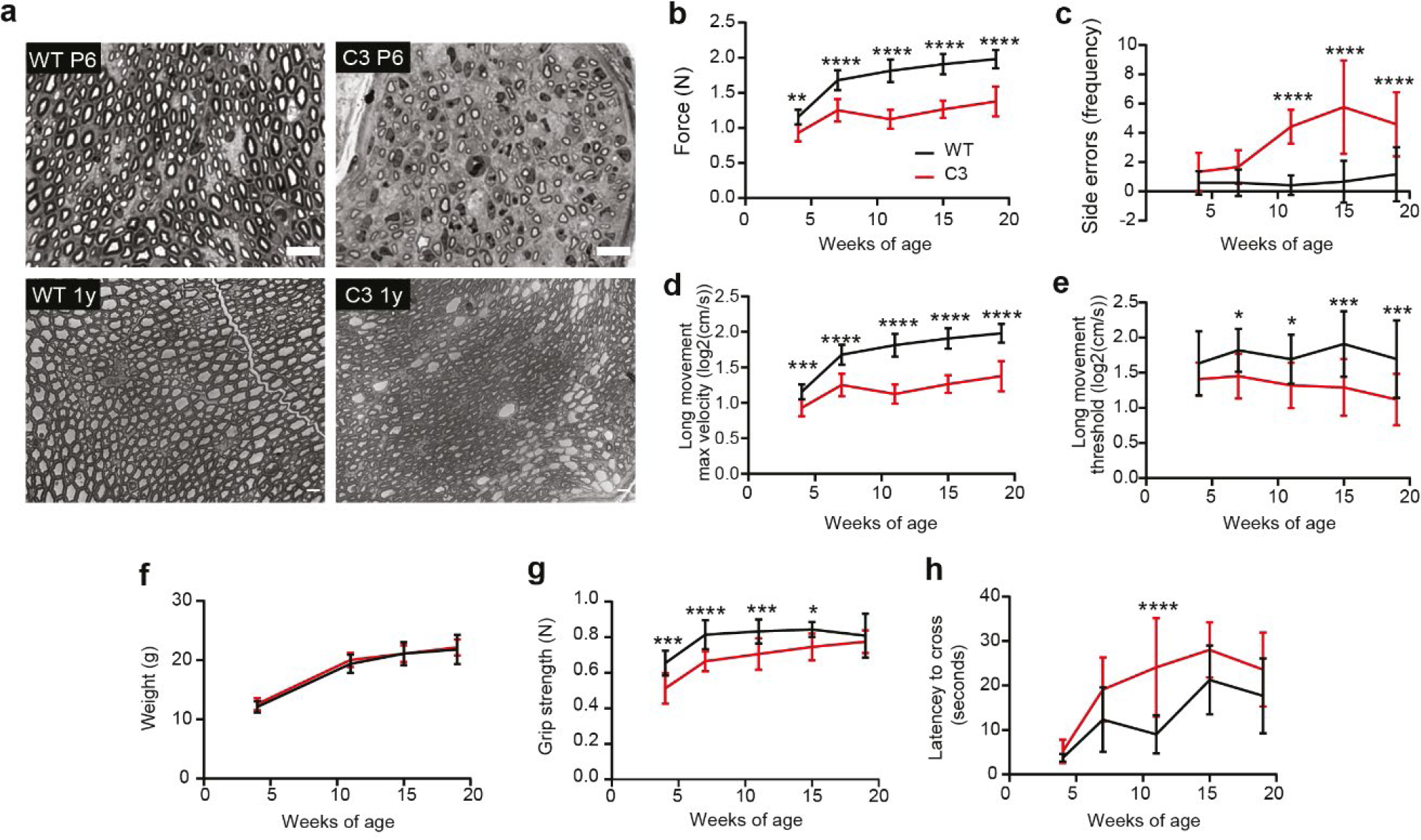
Functional follow-up of C3-PMP22 mice and wildtype littermate controls during development. **a)** Semi-thin cross-section from sciatic nerves of WT and C3 mice at postnatal day 6 and 1 year of age. Scale bar = 10 µm in all panels. **b-d)** Motor phenotype was assessed using grip strength (overall) (b), and also the number of errors while crossing a log beam were tracked (e). The maximal velocity long movement threshold (d) and the velocity long movement threshold (e) were monitored in the Phenotyper automated home cage. **f-h)** No difference in weight was recorded between C3 mice and their WT littermates at 3, 7, 12, 15, and 19 weeks old (f). In addition, the forepaw grip strength was significantly reduced in C3 mice (g), and also the latency to cross a beam was only significantly increased at 12 weeks of age in the C3 mice compared to age-matched WT controls (h). Statistical significance was evaluated in (b-h) with a two-way ANOVA test, followed by Fisher’s LSD test (* p < 0.05, ** p < 0.01, *** p < 0.001, **** p < 0.0001). Data are represented as the mean ± S.D., with 12 WT mice and 10-12 C3 mice per time point analyzed.

**Supplemental Figure 2.**
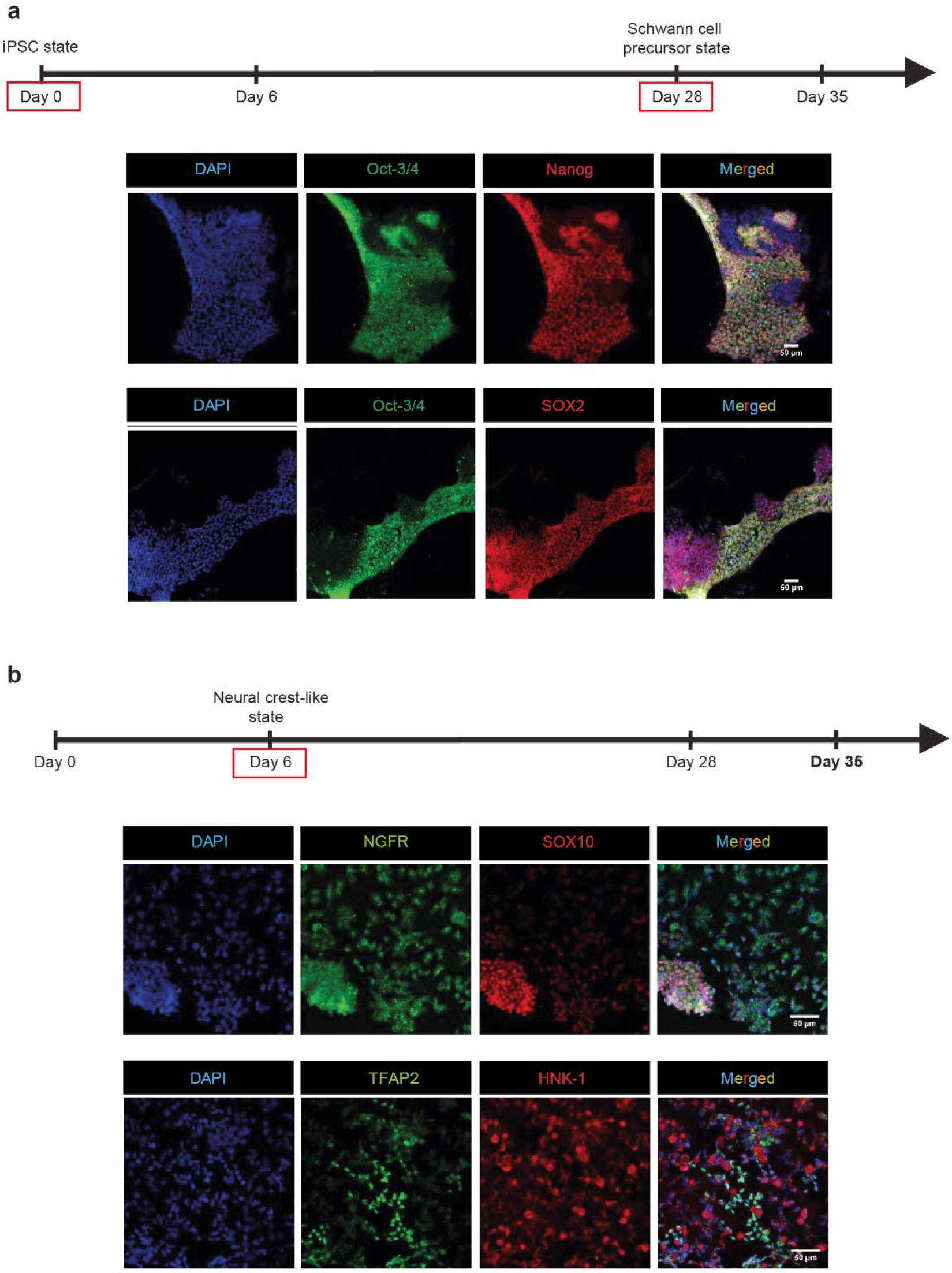
Expression of pluripotency and neural crest markers from days 0-6 of the iPSC-Schwann cell differentiation protocol. The control cell line from Sigma was used for the initial characterization of the iPSC-Schwann cell differentiation protocol. Their differentiation in the early stages of the protocol included validation of the expression of the a) pluripotency markers Oct-3/4, Nanog, and SOX2, and of the b) early neural crest markers, NGFR, TFAP2A, HNK-1, and SOX10 via immunocytochemistry. Nuclei (in blue) are visualized using DAPI. Scale bar is 50 µm for all images.

**Supplemental Figure 3.**
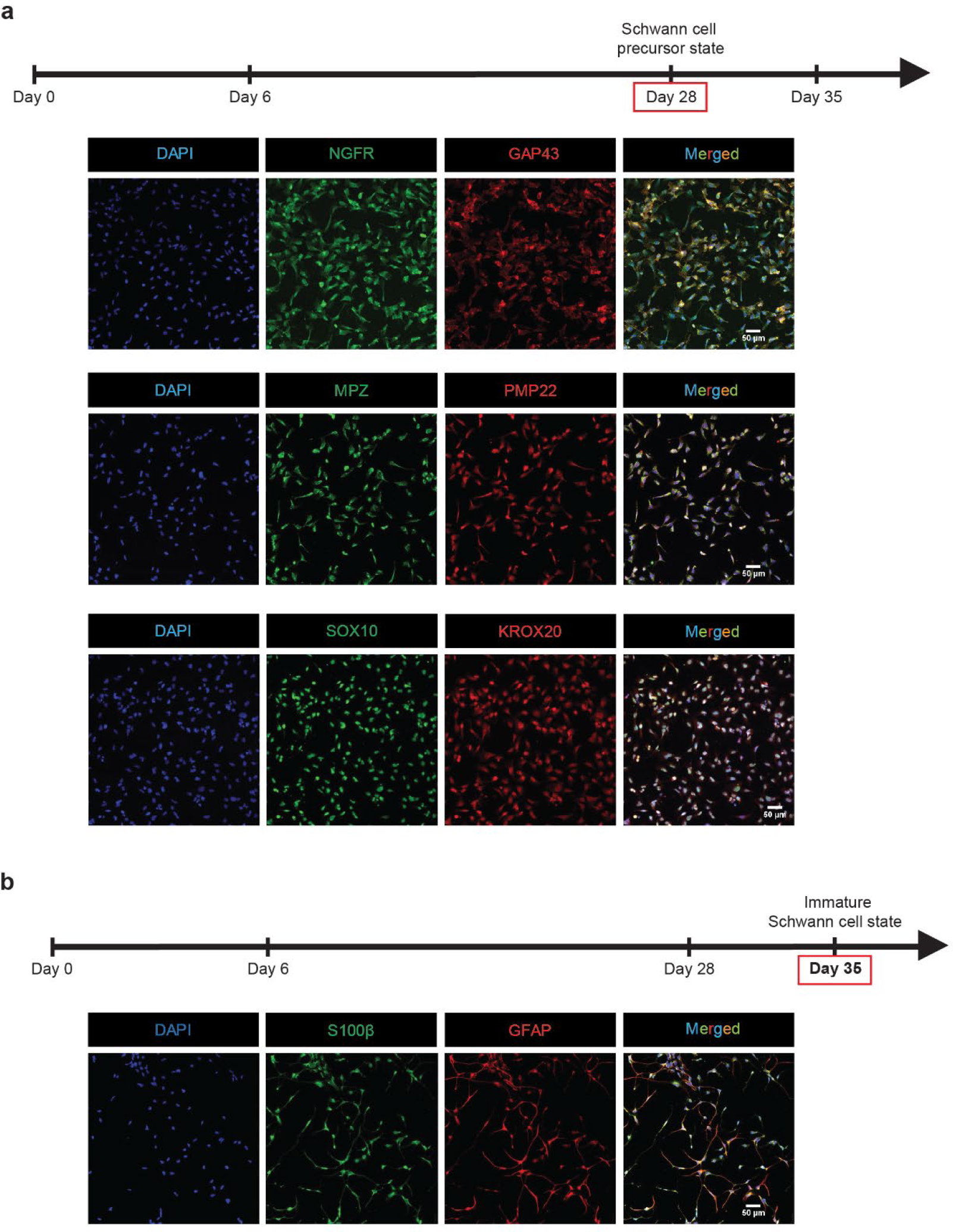
Expression of precursor and immature Schwann cell markers from days 28-35 of the iPSC-Schwann cell differentiation protocol. The control cell line from Sigma was used for the initial characterization of the iPSC-Schwann cell differentiation protocol. a) Their differentiation at the later stages of the protocol included expression analysis of Schwann cell precursor (SCP) markers such as NGFR, GAP43, MPZ (P0), PMP22, SOX10, and KROX20/EGRF2 on the protein level via immunofluorescence. b) At the final step, differentiated iPSCs are positive for the Schwann cell markers S100β and GFAP, and they have a bi-tripolar morphology. Nuclei (in blue) are visualized using DAPI. Scale bar is 50 µm in each image.

**Supplemental Figure 4.**
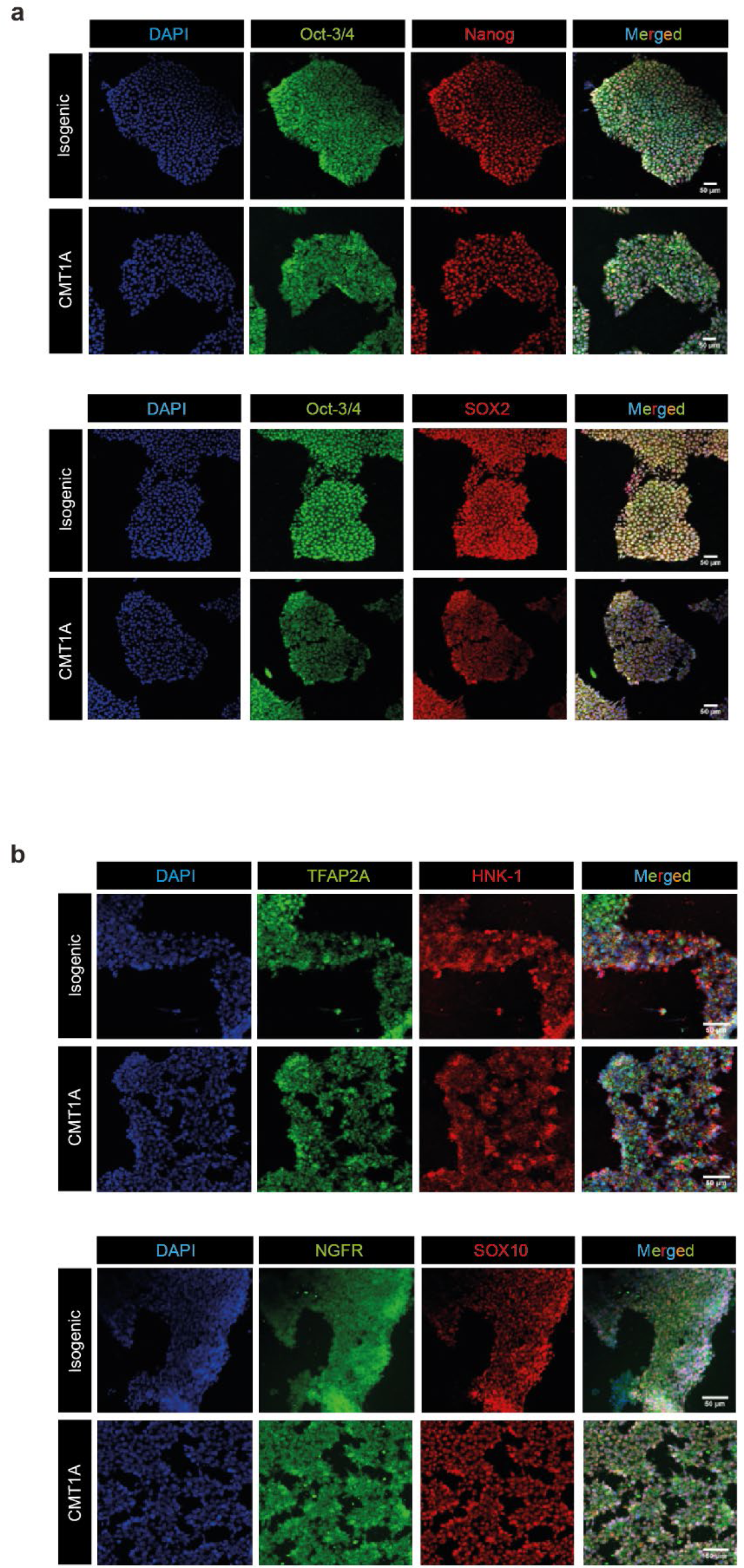
Expression of pluripotency markers of the CMT1A and isogenic iPSC line at the start of the differentiation protocol (iPSC-state) and after 6 days (neural crest level). a) Both the CMT1A and its isogenic control line are positive for several pluripotency markers, such as Nanog, Oct-3/4, and SOX2 at an iPSC level at the start of the differentiation. b) On day 6 of the protocol, the neural crest markers TFAP2A, HNK-1, NGFR, and SOX10 start to become expressed in both CMT1A and isogenic iPSCs. DAPI (in blue) was used as a nuclear counterstain. The scale bar is 50 µm in each image.

**Supplemental Figure 5.**
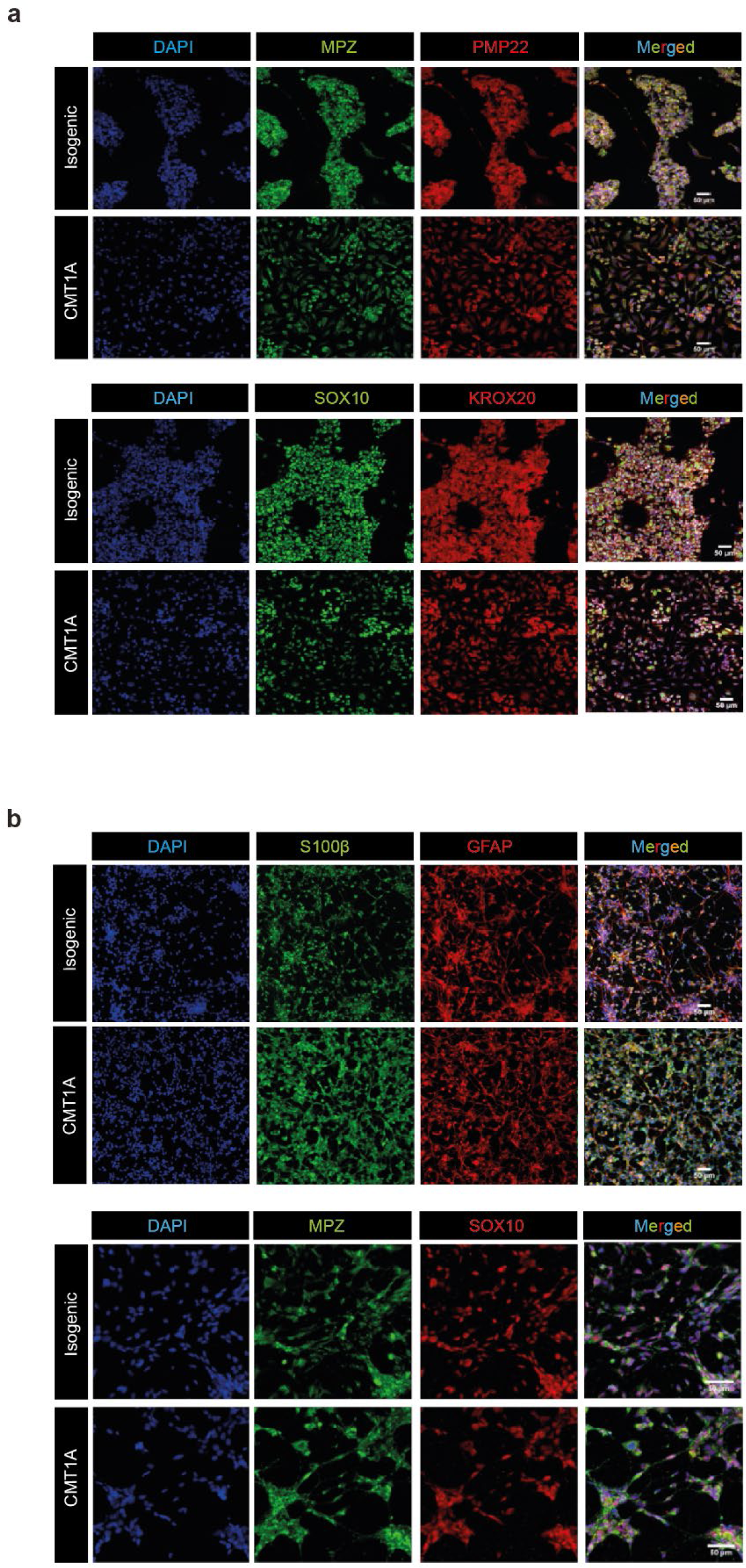
Expression of precursor and immature Schwann cell markers from day 28-35 of the iPSC-Schwann cell differentiation protocol for the CMT1A and its isogenic line. a) Both the CMT1A and its isogenic control line stained positive for MPZ (P0), PMP22, KROX20/EGRF2, and SOX10 on day 28 of the protocol. b) In addition, at day 35 of the differentiation protocol both the CMT1A and isogenic iPSC-SCPs stained positive for S100β, GFAP, MPZ (P0), and SOX10. DAPI (in blue) was used as a nuclear counterstain. The scale bar is 50 µm in each image.

**Supplemental Figure 6.**
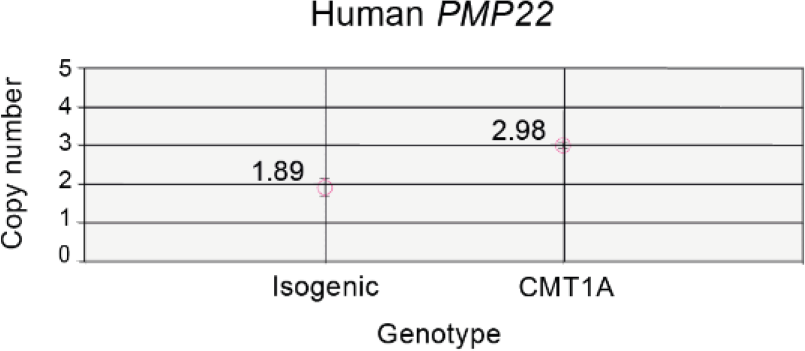
ddPCR analysis of isogenic and CMT1A iPSC-SCPs at 28 day of the Schwann cell differentiation protocol. Isogenic iPSC-SCPs express 1.89 copies of the human PMP22 and CMT1A iPSC-SCPs 2.98 copies, exactly one copy extra compared to the isogenic line.

**Supplemental Figure 7.**
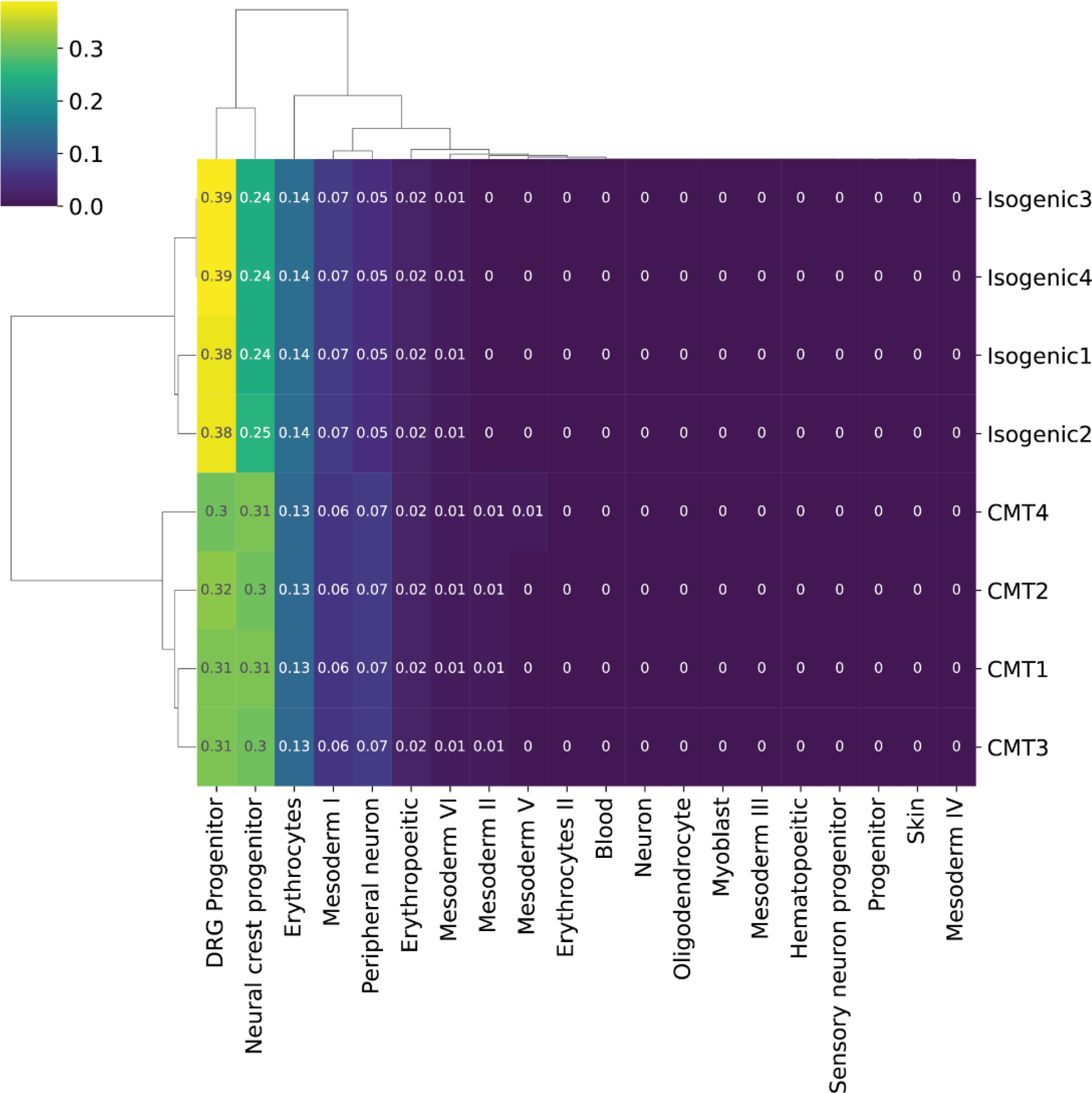
Bulk RNA-sequencing data maps to neural crest and DRG progenitor cells from a single cell atlas of a human developing spinal cord. DRG progenitors, which are defined as predominantly SOX10 positive, and neural crest progenitors, which are predominantly SOX10 and SOX2 positive cells, were the major cell types when mapped to the single cell atlas of the human developing spinal cord (Rayon *et al*., 2021). The term “Schwann cell precursor” was not used in the annotation of cell clusters in the study by Rayon *et al*., therefore it could not be used in the current cell cluster annotation in this figure.

**Supplemental Figure 8.**
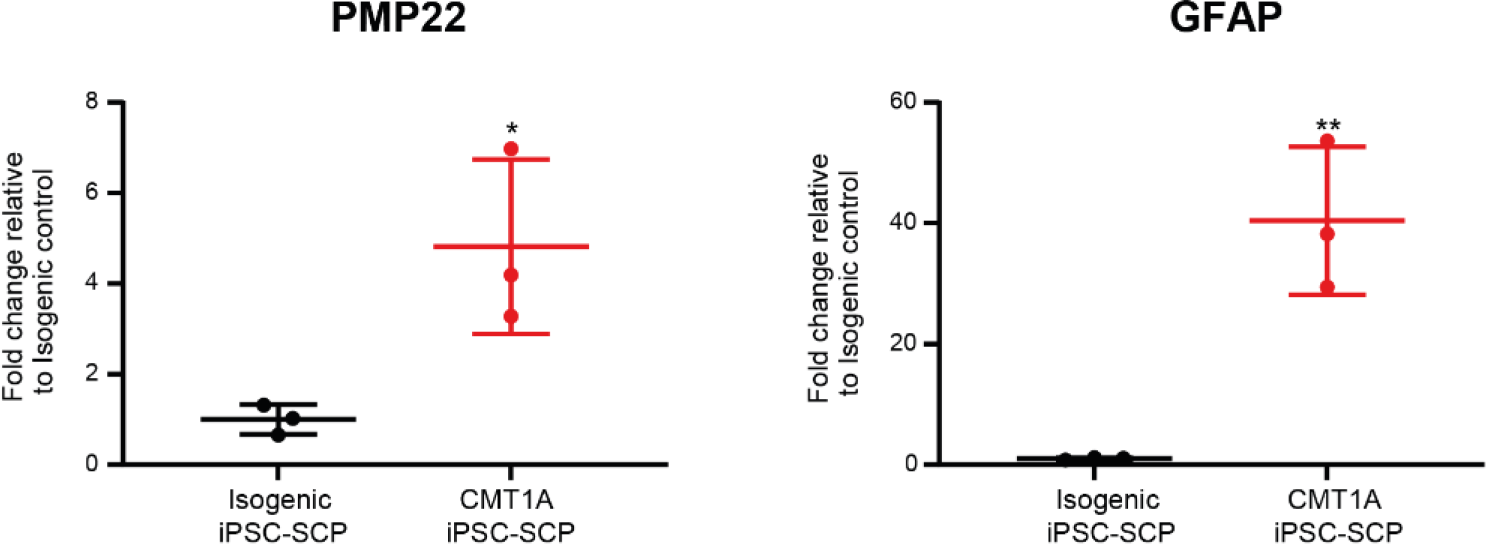
qPCR data indicates that PMP22 and GFAP mRNA are upregulated in CMT1A iPSC-SCPs at day 28 of the SC differentiation protocol when cultured in high BSA (0.5%) Data represent 3 independent differentiations, at independent time points. Data are represented as mean ± S.D. Statistical significance was determined using a two-tailed, unpaired t test (* p < 0.05, ** p < 0.01).

**Supplemental Figure 9.**
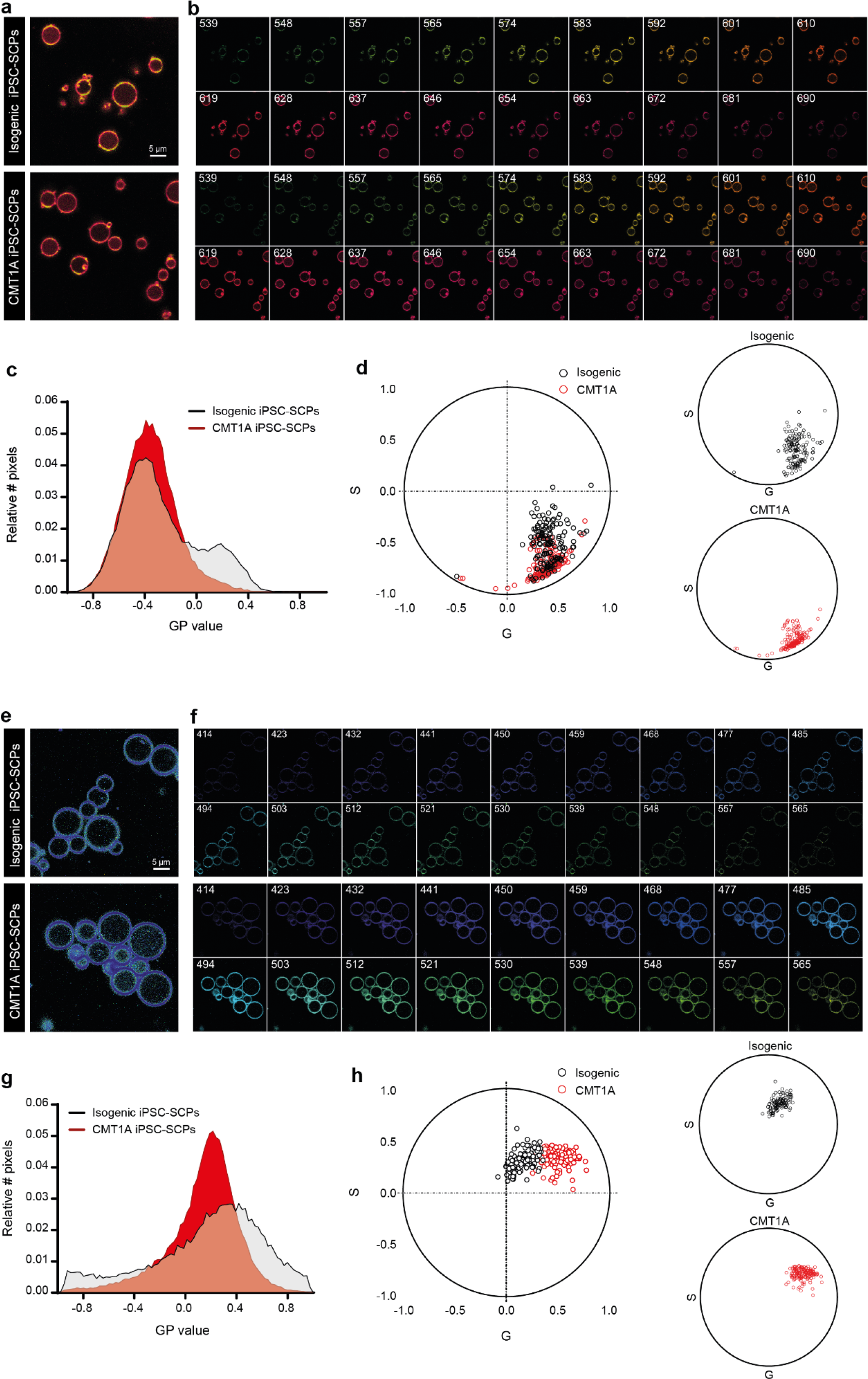
Spectral imaging with Di-4-ANEPPDHQ and Laurdan indicates decreased membrane order in the plasma membrane of CMT1A iPSC-SCPs. GPMVs were generated, and spectral (lambda, λ) confocal imaging was performed at different wavelengths with either Di-4-ANEPPDHQ or Laurdan as described in the methods. **a, b)** Photomicrographs of Di-4-ANEPPDHQ stained GPMVs from isogenic vs CMT1A iPSC-SCPs, including images with split wavelengths that are used to calculate the GPem = (IB(λ560) – IR(λ650)) / (IB(λ560) + IR(λ650)), ranging from -1 (liquid disordered, Ld) to +1 (liquid ordered, Lo). CMT1A: n = 164 and isogenic: n = 142 GPMVs analyzed, three independent experiments, data of one representative experiment are shown. c) Histogram indicating the distribution and amount of pixels per GP value for GPMVs from CMT1A (in red) and isogenic (in black) iPSC-SCPs labelled with Di-4-ANEPPDHQ. Histogram indicating the distribution and amount of pixels per GP value for Di-4-ANEPPDHQ GPMVs. **d)** Spectral phasor plots also indicate significant differences in data distribution between isogenic and CMT1A GPMVs (multivariate ANOVA analysis, p<0.0001, Eta Squared = 0.35). **e, f)** Photomicrographs of Laurdan stained GPMVs from isogenic vs CMT1A iPSC-SCPs, including images with split wavelengths that are used to calculate the GP = (IB(λ440) – IR(λ 490)) / (IB(λ 440) + IR(λ 490)). CMT1A: n = 151 and isogenic: n = 112 GPMVs analyzed, two independent experiments, data of one representative experiment are shown. **g)** Histogram indicating the distribution and amount of pixels per GP value for GPMVs from CMT1A (in red) and isogenic (in black) iPSC-SCPs labelled with Laurdan. **h)** Spectral phasor plots also indicate significant differences in data distribution between isogenic and CMT1A GPMVs (multivariate ANOVA analysis, p<0.0001, Eta Squared = 0.73).

**Supplemental Figure 10.**
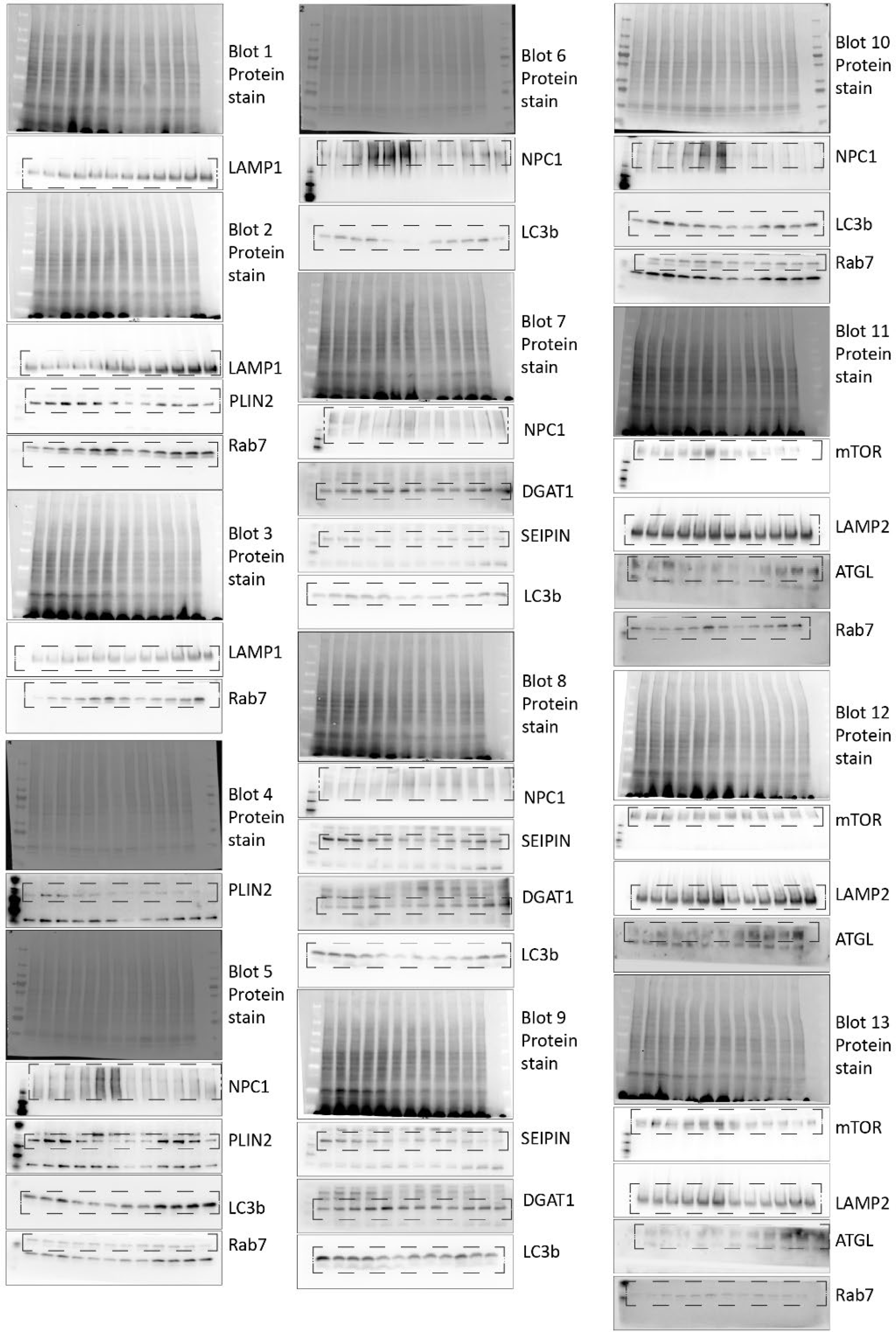
Overview of Western blots used for quantifications in Fig. 6 of the main text. 5 experiments with markers stained at least 3 times for quantifications. After protein stain, markers stained for each blot are highlighted. Blots are not ordered in a sequential manner.

## Notes

### Competing Interest Statement

The authors have declared no competing interest.

## References

Amici, S.A., Dunn, W.A., Jr., and Notterpek, L. (2007). Developmental abnormalities in the nerves of peripheral myelin protein 22-deficient mice. J Neurosci Res 85, 238–249. 10.1002/jnr.21118.

Aron, M., Browning, R., Carugo, D., Sezgin, E., Bernardino de la Serna, J., Eggeling, C., and Stride, E. (2017). Spectral imaging toolbox: segmentation, hyperstack reconstruction, and batch processing of spectral images for the determination of cell and model membrane lipid order. BMC Bioinformatics 18, 254. 10.1186/s12859-017-1656-2.

Bademosi, A.T., Lauwers, E., Padmanabhan, P., Odierna, L., Chai, Y.J., Papadopulos, A., Goodhill, G.J., Verstreken, P., van Swinderen, B., and Meunier, F.A. (2017). In vivo single-molecule imaging of syntaxin1A reveals polyphosphoinositide- and activity-dependent trapping in presynaptic nanoclusters. Nat Commun 8, 13660. 10.1038/ncomms13660.

Benoy, V., Van Helleputte, L., Prior, R., d’Ydewalle, C., Haeck, W., Geens, N., Scheveneels, W., Schevenels, B., Cader, M.Z., Talbot, K., et al. (2018). HDAC6 is a therapeutic target in mutant GARS-induced Charcot-Marie-Tooth disease. Brain 141, 673–687. 10.1093/brain/awx375.

Bhat, M.A. (2003). Molecular organization of axo-glial junctions. Curr Opin Neurobiol 13, 552–559. 10.1016/j.conb.2003.09.004.

Blanchette-Mackie, E.J., Dwyer, N.K., Barber, T., Coxey, R.A., Takeda, T., Rondinone, C.M., Theodorakis, J.L., Greenberg, A.S., and Londos, C. (1995). Perilipin is located on the surface layer of intracellular lipid droplets in adipocytes. J Lipid Res 36, 1211–1226.

Blom, T.S., Linder, M.D., Snow, K., Pihko, H., Hess, M.W., Jokitalo, E., Veckman, V., Syvanen, A.C., and Ikonen, E. (2003). Defective endocytic trafficking of NPC1 and NPC2 underlying infantile Niemann-Pick type C disease. Hum Mol Genet 12, 257–272. 10.1093/hmg/ddg025.

Bosse, F., Brodbeck, J., and Muller, H.W. (1999). Post-transcriptional regulation of the peripheral myelin protein gene PMP22/gas3. J Neurosci Res 55, 164–177. 10.1002/(SICI)1097-4547(19990115)55:2<164::AID-JNR4>3.0.CO;2-9.

Brady, S.T., Witt, A.S., Kirkpatrick, L.L., de Waegh, S.M., Readhead, C., Tu, P.H., and Lee, V.M. (1999). Formation of compact myelin is required for maturation of the axonal cytoskeleton. J Neurosci 19, 7278–7288. 10.1523/JNEUROSCI.19-17-07278.1999.

Brewer, J., Bernardino de la Serna, J., Wagner, K., and Bagatolli, L.A. (2010). Multiphoton excitation fluorescence microscopy in planar membrane systems. Biochim Biophys Acta 1798, 1301–1308. 10.1016/j.bbamem.2010.02.024.

Browning, R.J., Aron, M., Booth, A., Rademeyer, P., Wing, S., Brans, V., Shrivastava, S., Carugo, D., and Stride, E. (2020). Spectral Imaging for Microbubble Characterization. Langmuir 36, 609–617. 10.1021/acs.langmuir.9b03828.

Byravan, S., and Campagnoni, A.T. (1994). Serum factors and hydrocortisone influence the synthesis of myelin basic proteins in mouse brain primary cu--ltures. Int J Dev Neurosci 12, 343–351. 10.1016/0736-5748(94)90084-1.

Cartwright, B.R., Binns, D.D., Hilton, C.L., Han, S., Gao, Q., and Goodman, J.M. (2015). Seipin performs dissectible functions in promoting lipid droplet biogenesis and regulating droplet morphology. Mol Biol Cell 26, 726–739. 10.1091/mbc.E14-08-1303.

Chang, T. Y., C. C. Chang, N. Ohgami & Y. Yamauchi (2006) Cholesterol sensing, trafficking, and esterification. Annu Rev Cell Dev Biol, 22, 129–57.

Chittoor, V.G., Sooyeon, L., Rangaraju, S., Nicks, J.R., Schmidt, J.T., Madorsky, I., Narvaez, D.C., and Notterpek, L. (2013). Biochemical characterization of protein quality control mechanisms during disease progression in the C22 mouse model of CMT1A. ASN Neuro 5, e00128. 10.1042/AN20130024.

Christenson, L.K., and Devoto, L. (2003). Cholesterol transport and steroidogenesis by the corpus luteum. Reprod Biol Endocrinol 1, 90. 10.1186/1477-7827-1-90.

Crane, J.M., and Tamm, L.K. (2004). Role of cholesterol in the formation and nature of lipid rafts in planar and spherical model membranes. Biophys J 86, 2965–2979. 10.1016/S0006-3495(04)74347-7.

Desarnaud, F., Do Thi, A.N., Brown, A.M., Lemke, G., Suter, U., Baulieu, E.E., and Schumacher, M. (1998). Progesterone stimulates the activity of the promoters of peripheral myelin protein-22 and protein zero genes in Schwann cells. J Neurochem 71, 1765–1768. 10.1046/j.1471-4159.1998.71041765.x.

Duncan, R.E., Ahmadian, M., Jaworski, K., Sarkadi-Nagy, E., and Sul, H.S. (2007). Regulation of lipolysis in adipocytes. Annu Rev Nutr 27, 79–101. 10.1146/annurev.nutr.27.061406.093734.

Fledrich, R., Abdelaal, T., Rasch, L., Bansal, V., Schutza, V., Brugger, B., Luchtenborg, C., Prukop, T., Stenzel, J., Rahman, R.U., et al. (2018). Targeting myelin lipid metabolism as a potential therapeutic strategy in a model of CMT1A neuropathy. Nat Commun 9, 3025. 10.1038/s41467-018-05420-0.

Fledrich, R., Schlotter-Weigel, B., Schnizer, T.J., Wichert, S.P., Stassart, R.M., Meyer zu Horste, G., Klink, A., Weiss, B.G., Haag, U., Walter, M.C., et al. (2012). A rat model of Charcot-Marie-Tooth disease 1A recapitulates disease variability and supplies biomarkers of axonal loss in patients. Brain 135, 72–87. 10.1093/brain/awr322.

Fledrich, R., Stassart, R.M., Klink, A., Rasch, L.M., Prukop, T., Haag, L., Czesnik, D., Kungl, T., Abdelaal, T.A., Keric, N., et al. (2014). Soluble neuregulin-1 modulates disease pathogenesis in rodent models of Charcot-Marie-Tooth disease 1A. Nat Med 20, 1055–1061. 10.1038/nm.3664.

Fortun, J., Go, J.C., Li, J., Amici, S.A., Dunn, W.A., Jr., and Notterpek, L. (2006). Alterations in degradative pathways and protein aggregation in a neuropathy model based on PMP22 overexpression. Neurobiol Dis 22, 153–164. 10.1016/j.nbd.2005.10.010.

Frolov, A., Zielinski, S.E., Crowley, J.R., Dudley-Rucker, N., Schaffer, J.E., and Ory, D.S. (2003). NPC1 and NPC2 regulate cellular cholesterol homeostasis through generation of low density lipoprotein cholesterol-derived oxysterols. J Biol Chem 278, 25517–25525. 10.1074/jbc.M302588200.

Gomis-Coloma, C., Velasco-Aviles, S., Gomez-Sanchez, J.A., Casillas-Bajo, A., Backs, J., and Cabedo, H. (2018). Class IIa histone deacetylases link cAMP signaling to the myelin transcriptional program of Schwann cells. J Cell Biol 217, 1249–1268. 10.1083/jcb.201611150.

Grisan, F., Iannucci, L.F., Surdo, N.C., Gerbino, A., Zanin, S., Di Benedetto, G., Pozzan, T., and Lefkimmiatis, K. (2021). PKA compartmentalization links cAMP signaling and autophagy. Cell Death Differ 28, 2436–2449. 10.1038/s41418-021-00761-8.

Guo, W., Naujock, M., Fumagalli, L., Vandoorne, T., Baatsen, P., Boon, R., Ordovas, L., Patel, A., Welters, M., Vanwelden, T., et al. (2017). HDAC6 inhibition reverses axonal transport defects in motor neurons derived from FUS-ALS patients. Nat Commun 8, 861. 10.1038/s41467-017-00911-y.

Hanemann, C.O., Stoll, G., D’Urso, D., Fricke, W., Martin, J.J., Van Broeckhoven, C., Mancardi, G.L., Bartke, I., and Muller, H.W. (1994). Peripheral myelin protein-22 expression in Charcot-Marie-Tooth disease type 1a sural nerve biopsies. J Neurosci Res 37, 654–659. 10.1002/jnr.490370513.

Harayama, T., and Shimizu, T. (2020). Roles of polyunsaturated fatty acids, from mediators to membranes. J Lipid Res 61, 1150–1160. 10.1194/jlr.R120000800.

Henne, W.M., Reese, M.L., and Goodman, J.M. (2018). The assembly of lipid droplets and their roles in challenged cells. EMBO J 37. 10.15252/embj.201898947.

Hoffmann, K., Nirmalananthan-Budau, N., and Resch-Genger, U. (2020). Fluorescence calibration standards made from broadband emitters encapsulated in polymer beads for fluorescence microscopy and flow cytometry. Anal Bioanal Chem 412, 6499–6507. 10.1007/s00216-020-02664-y.

Huxley, C., Passage, E., Manson, A., Putzu, G., Figarella-Branger, D., Pellissier, J.F., and Fontes, M. (1996). Construction of a mouse model of Charcot-Marie-Tooth disease type 1A by pronuclear injection of human YAC DNA. Hum Mol Genet 5, 563–569. 10.1093/hmg/5.5.563.

Jay, A.G., and Hamilton, J.A. (2017). Disorder Amidst Membrane Order: Standardizing Laurdan Generalized Polarization and Membrane Fluidity Terms. J Fluoresc 27, 243–249. 10.1007/s10895-016-1951-8.

Jin, L., Millard, A.C., Wuskell, J.P., Clark, H.A., and Loew, L.M. (2005). Cholesterol-enriched lipid domains can be visualized by di-4-ANEPPDHQ with linear and nonlinear optics. Biophys J 89, L04–06. 10.1529/biophysj.105.064816.

Katona, I., Wu, X., Feely, S.M., Sottile, S., Siskind, C.E., Miller, L.J., Shy, M.E., and Li, J. (2009). PMP22 expression in dermal nerve myelin from patients with CMT1A. Brain 132, 1734–1740. 10.1093/brain/awp113.

Kechkar, A., Nair, D., Heilemann, M., Choquet, D., and Sibarita, J.B. (2013). Real-time analysis and visualization for single-molecule based super-resolution microscopy. PLoS One 8, e62918. 10.1371/journal.pone.0062918.

Kim, H.S., Lee, J., Lee, D.Y., Kim, Y.D., Kim, J.Y., Lim, H.J., Lim, S., and Cho, Y.S. (2017). Schwann Cell Precursors from Human Pluripotent Stem Cells as a Potential Therapeutic Target for Myelin Repair. Stem Cell Reports 8, 1714–1726. 10.1016/j.stemcr.2017.04.011.

Krauter, D.E., D.; Hartmann, T. J.; Volkmann, S.; Kungl, T.; Fledrich, R.; Goebbels, S.; Nave, K-A.; Sereda M. W. (2021). Inversely proportional myelin growth due to altered Pmp22 gene dosage identifies PI3K/Akt/mTOR signaling as a novel therapeutic target in HNPP. bioRxiv. 10.1101/2021.11.08.467756.

Kurat, C.F., Wolinski, H., Petschnigg, J., Kaluarachchi, S., Andrews, B., Natter, K., and Kohlwein, S.D. (2009). Cdk1/Cdc28-dependent activation of the major triacylglycerol lipase Tgl4 in yeast links lipolysis to cell-cycle progression. Mol Cell 33, 53–63. 10.1016/j.molcel.2008.12.019.

Le Douarin, N., Dulac, C., Dupin, E., and Cameron-Curry, P. (1991). Glial cell lineages in the neural crest. Glia 4, 175–184. 10.1002/glia.440040209.

Lee, S., Amici, S., Tavori, H., Zeng, W.M., Freeland, S., Fazio, S., and Notterpek, L. (2014). PMP22 is critical for actin-mediated cellular functions and for establishing lipid rafts. J Neurosci 34, 16140–16152. 10.1523/JNEUROSCI.1908-14.2014.

Liebisch, G., Fahy, E., Aoki, J., Dennis, E.A., Durand, T., Ejsing, C.S., Fedorova, M., Feussner, I., Griffiths, W.J., Kofeler, H., et al. (2020). Update on LIPID MAPS classification, nomenclature, and shorthand notation for MS-derived lipid structures. J Lipid Res 61, 1539–1555. 10.1194/jlr.S120001025.

Liebisch, G., Vizcaino, J.A., Kofeler, H., Trotzmuller, M., Griffiths, W.J., Schmitz, G., Spener, F., and Wakelam, M.J.O. (2013). Shorthand notation for lipid structures derived from mass spectrometry. J Lipid Res 54, 1523–1530. 10.1194/jlr.M033506.

Lingwood, D., and Simons, K. (2010). Lipid rafts as a membrane-organizing principle. Science 327, 46–50. 10.1126/science.1174621.

Liu, B., Xin, W., Tan, J.R., Zhu, R.P., Li, T., Wang, D., Kan, S.S., Xiong, D.K., Li, H.H., Zhang, M.M., et al. (2019). Myelin sheath structure and regeneration in peripheral nerve injury repair. Proc Natl Acad Sci U S A 116, 22347–22352. 10.1073/pnas.1910292116.

Malacrida, L., Astrada, S., Briva, A., Bollati-Fogolin, M., Gratton, E., and Bagatolli, L.A. (2016). Spectral phasor analysis of LAURDAN fluorescence in live A549 lung cells to study the hydration and time evolution of intracellular lamellar body-like structures. Biochim Biophys Acta 1858, 2625–2635. 10.1016/j.bbamem.2016.07.017.

Malacrida, L., Gratton, E., and Jameson, D.M. (2015). Model-free methods to study membrane environmental probes: a comparison of the spectral phasor and generalized polarization approaches. Methods Appl Fluoresc 3. 10.1088/2050-6120/3/4/047001.

Marinko, J.T., Kenworthy, A.K., and Sanders, C.R. (2020). Peripheral myelin protein 22 preferentially partitions into ordered phase membrane domains. Proc Natl Acad Sci U S A 117, 14168–14177. 10.1073/pnas.2000508117.

Martini, R. (2001). The effect of myelinating Schwann cells on axons. Muscle Nerve 24, 456–466. 10.1002/mus.1027.

Metherall, J.E., Waugh, K., and Li, H. (1996). Progesterone inhibits cholesterol biosynthesis in cultured cells. Accumulation of cholesterol precursors. J Biol Chem 271, 2627–2633. 10.1074/jbc.271.5.2627.

Mirsky, R., Jessen, K.R., Brennan, A., Parkinson, D., Dong, Z., Meier, C., Parmantier, E., and Lawson, D. (2002). Schwann cells as regulators of nerve development. J Physiol Paris 96, 17–24. 10.1016/s0928-4257(01)00076-6.

Mittendorf, K.F., Marinko, J.T., Hampton, C.M., Ke, Z., Hadziselimovic, A., Schlebach, J.P., Law, C.L., Li, J., Wright, E.R., Sanders, C.R., and Ohi, M.D. (2017). Peripheral myelin protein 22 alters membrane architecture. Sci Adv 3, e1700220. 10.1126/sciadv.1700220.

Morgan, L., Jessen, K.R., and Mirsky, R. (1991). The effects of cAMP on differentiation of cultured Schwann cells: progression from an early phenotype (04+) to a myelin phenotype (P0+, GFAP-, N-CAM-, NGF-receptor-) depends on growth inhibition. J Cell Biol 112, 457–467. 10.1083/jcb.112.3.457.

Mukherjee-Clavin, B., Mi, R., Kern, B., Choi, I.Y., Lim, H., Oh, Y., Lannon, B., Kim, K.J., Bell, S., Hur, J.K., et al. (2019). Comparison of three congruent patient-specific cell types for the modelling of a human genetic Schwann-cell disorder. Nat Biomed Eng 3, 571–582. 10.1038/s41551-019-0381-8.

Olzmann, J.A., and Carvalho, P. (2019). Dynamics and functions of lipid droplets. Nat Rev Mol Cell Biol 20, 137–155. 10.1038/s41580-018-0085-z.

Owen, D.M., Williamson, D., Magenau, A., and Gaus, K. (2012). Optical techniques for imaging membrane domains in live cells (live-cell palm of protein clustering). Methods Enzymol 504, 221–235. 10.1016/B978-0-12-391857-4.00011-2.

Pang, Z., Chong, J., Zhou, G., de Lima Morais, D.A., Chang, L., Barrette, M., Gauthier, C., Jacques, P.E., Li, S., and Xia, J. (2021). MetaboAnalyst 5.0: narrowing the gap between raw spectra and functional insights. Nucleic Acids Res 49, W388–W396. 10.1093/nar/gkab382.

Parasassi, T., and Gratton, E. (1995). Membrane lipid domains and dynamics as detected by Laurdan fluorescence. J Fluoresc 5, 59–69. 10.1007/BF00718783.

Parisio, G., Marini, A., Biancardi, A., Ferrarini, A., and Mennucci, B. (2011). Polarity-sensitive fluorescent probes in lipid bilayers: bridging spectroscopic behavior and microenvironment properties. J Phys Chem B 115, 9980–9989. 10.1021/jp205163w.

Pertusa, M., Morenilla-Palao, C., Carteron, C., Viana, F., and Cabedo, H. (2007). Transcriptional control of cholesterol biosynthesis in Schwann cells by axonal neuregulin 1. J Biol Chem 282, 28768–28778. 10.1074/jbc.M701878200.

Prior, R., Verschoren, S., Vints, K., Jaspers, T., Rossaert, E., Klingl, Y.E., Silva, A., Hersmus, N., Van Damme, P., and Van Den Bosch, L. (2022). HDAC3 Inhibition Stimulates Myelination in a CMT1A Mouse Model. Mol Neurobiol 59, 3414–3430. 10.1007/s12035-022-02782-x.

Raeymaekers, P., Timmerman, V., Nelis, E., De Jonghe, P., Hoogendijk, J.E., Baas, F., Barker, D.F., Martin, J.J., De Visser, M., Bolhuis, P.A., and, et al. (1991). Duplication in chromosome 17p11.2 in Charcot-Marie-Tooth neuropathy type 1a (CMT 1a). The HMSN Collaborative Research Group. Neuromuscul Disord 1, 93–97. 10.1016/0960-8966(91)90055-w.

Rambold, A.S., Cohen, S., and Lippincott-Schwartz, J. (2015). Fatty acid trafficking in starved cells: regulation by lipid droplet lipolysis, autophagy, and mitochondrial fusion dynamics. Dev Cell 32, 678–692. 10.1016/j.devcel.2015.01.029.

Ravelo, K.M., Andersen, N.D., and Monje, P.V. (2018). Magnetic-Activated Cell Sorting for the Fast and Efficient Separation of Human and Rodent Schwann Cells from Mixed Cell Populations. Methods Mol Biol 1739, 87–109. 10.1007/978-1-4939-7649-2_6.

Rayon, T., Maizels, R.J., Barrington, C., and Briscoe, J. (2021). Single-cell transcriptome profiling of the human developing spinal cord reveals a conserved genetic programme with human-specific features. Development 148. 10.1242/dev.199711.

Redondo-Morata, L., Giannotti, M.I., and Sanz, F. (2012). Influence of cholesterol on the phase transition of lipid bilayers: a temperature-controlled force spectroscopy study. Langmuir 28, 12851–12860. 10.1021/la302620t.

Richner, M., Jager, S.B., Siupka, P., and Vaegter, C.B. (2017). Hydraulic Extrusion of the Spinal Cord and Isolation of Dorsal Root Ganglia in Rodents. J Vis Exp. 10.3791/55226.

Robaglia-Schlupp, A., Pizant, J., Norreel, J.C., Passage, E., Saberan-Djoneidi, D., Ansaldi, J.L., Vinay, L., Figarella-Branger, D., Levy, N., Clarac, F., et al. (2002). PMP22 overexpression causes dysmyelination in mice. Brain 125, 2213–2221. 10.1093/brain/awf230.

Robertson, A.M., Huxley, C., King, R.H., and Thomas, P.K. (1999). Development of early postnatal peripheral nerve abnormalities in Trembler-J and PMP22 transgenic mice. J Anat 195 ( *Pt 3*), 331–339. 10.1046/j.1469-7580.1999.19530331.x.

Saher, G., and Stumpf, S.K. (2015). Cholesterol in myelin biogenesis and hypomyelinating disorders. Biochim Biophys Acta 1851, 1083–1094. 10.1016/j.bbalip.2015.02.010.

Salzer, J.L., and Zalc, B. (2016). Myelination. Curr Biol 26, R971–R975. 10.1016/j.cub.2016.07.074.

Sanchez, S.A., Tricerri, M.A., and Gratton, E. (2012). Laurdan generalized polarization fluctuations measures membrane packing micro-heterogeneity in vivo. Proc Natl Acad Sci U S A 109, 7314–7319. 10.1073/pnas.1118288109.

Sereda, M., Griffiths, I., Puhlhofer, A., Stewart, H., Rossner, M.J., Zimmerman, F., Magyar, J.P., Schneider, A., Hund, E., Meinck, H.M., et al. (1996). A transgenic rat model of Charcot-Marie-Tooth disease. Neuron 16, 1049–1060. 10.1016/s0896-6273(00)80128-2.

Sereda, M.W., Meyer zu Horste, G., Suter, U., Uzma, N., and Nave, K.A. (2003). Therapeutic administration of progesterone antagonist in a model of Charcot-Marie-Tooth disease (CMT-1A). Nat Med 9, 1533–1537. 10.1038/nm957.

Sezgin, E., Kaiser, H.J., Baumgart, T., Schwille, P., Simons, K., and Levental, I. (2012). Elucidating membrane structure and protein behavior using giant plasma membrane vesicles. Nat Protoc 7, 1042–1051. 10.1038/nprot.2012.059.

Sezgin, E., Levental, I., Mayor, S., and Eggeling, C. (2017). The mystery of membrane organization: composition, regulation and roles of lipid rafts. Nat Rev Mol Cell Biol 18, 361–374. 10.1038/nrm.2017.16.

Sezgin, E., Waithe, D., Bernardino de la Serna, J., and Eggeling, C. (2015). Spectral imaging to measure heterogeneity in membrane lipid packing. Chemphyschem 16, 1387–1394. 10.1002/cphc.201402794.

Shi, L., Huang, L., He, R., Huang, W., Wang, H., Lai, X., Zou, Z., Sun, J., Ke, Q., Zheng, M., et al. (2018). Modeling the Pathogenesis of Charcot-Marie-Tooth Disease Type 1A Using Patient-Specific iPSCs. Stem Cell Reports 10, 120–133. 10.1016/j.stemcr.2017.11.013.

Siems, S.B., Jahn, O., Eichel, M.A., Kannaiyan, N., Wu, L.M.N., Sherman, D.L., Kusch, K., Hesse, D., Jung, R.B., Fledrich, R., et al. (2020). Proteome profile of peripheral myelin in healthy mice and in a neuropathy model. Elife 9. 10.7554/eLife.51406.

Simons, K., and Ehehalt, R. (2002). Cholesterol, lipid rafts, and disease. J Clin Invest 110, 597–603. 10.1172/JCI16390.

Simons, K., and Toomre, D. (2000). Lipid rafts and signal transduction. Nat Rev Mol Cell Biol 1, 31–39. 10.1038/35036052.

Slenders, E., Seneca, S., Pramanik, S.K., Smisdom, N., Adriaensens, P., vandeVen, M., Ethirajan, A., and Ameloot, M. (2018). Dynamics of the phospholipid shell of microbubbles: a fluorescence photoselection and spectral phasor approach. Chem Commun (Camb) 54, 4854–4857. 10.1039/c8cc01012a.

Socas, L.B.P., and Ambroggio, E.E. (2022). Introducing the multi-dimensional spectral phasors: a tool for the analysis of fluorescence excitation-emission matrices. Methods Appl Fluoresc 10. 10.1088/2050-6120/ac5389.

Stassart, R.M., and Woodhoo, A. (2021). Axo-glial interaction in the injured PNS. Dev Neurobiol 81, 490–506. 10.1002/dneu.22771.

Stefl, M., James, N.G., Ross, J.A., and Jameson, D.M. (2011). Applications of phasors to in vitro time-resolved fluorescence measurements. Anal Biochem 410, 62–69. 10.1016/j.ab.2010.11.010.

Stoklund Dittlau, K., Krasnow, E.N., Fumagalli, L., Vandoorne, T., Baatsen, P., Kerstens, A., Giacomazzi, G., Pavie, B., Rossaert, E., Beckers, J., et al. (2021). Human motor units in microfluidic devices are impaired by FUS mutations and improved by HDAC6 inhibition. Stem Cell Reports 16, 2213–2227. 10.1016/j.stemcr.2021.03.029.

Subramanian, K., and Balch, W.E. (2008). NPC1/NPC2 function as a tag team duo to mobilize cholesterol. Proc Natl Acad Sci U S A 105, 15223–15224. 10.1073/pnas.0808256105.

Suhaj, A., Gowland, D., Bonini, N., Owen, D.M., and Lorenz, C.D. (2020). Laurdan and Di-4-ANEPPDHQ Influence the Properties of Lipid Membranes: A Classical Molecular Dynamics and Fluorescence Study. J Phys Chem B 124, 11419–11430. 10.1021/acs.jpcb.0c09496.

Sztalryd, C., and Brasaemle, D.L. (2017). The perilipin family of lipid droplet proteins: Gatekeepers of intracellular lipolysis. Biochim Biophys Acta Mol Cell Biol Lipids 1862, 1221–1232. 10.1016/j.bbalip.2017.07.009.

Taniguchi, M., and Okazaki, T. (2020). Ceramide/Sphingomyelin Rheostat Regulated by Sphingomyelin Synthases and Chronic Diseases in Murine Models. J Lipid Atheroscler 9, 380–405. 10.12997/jla.2020.9.3.380.

Timmerman, V., Nelis, E., Van Hul, W., Nieuwenhuijsen, B.W., Chen, K.L., Wang, S., Ben Othman, K., Cullen, B., Leach, R.J., Hanemann, C.O., and, et al. (1992). The peripheral myelin protein gene PMP-22 is contained within the Charcot-Marie-Tooth disease type 1A duplication. Nat Genet 1, 171–175. 10.1038/ng0692-171.

Ugland, H., Naderi, S., Brech, A., Collas, P., and Blomhoff, H.K. (2011). cAMP induces autophagy via a novel pathway involving ERK, cyclin E and Beclin 1. Autophagy 7, 1199–1211. 10.4161/auto.7.10.16649.

Valmassoi, J., and Schwarz, R.I. (1988). High serum levels interfere with the normal differentiated state of avian tendon cells by altering translational regulation. Exp Cell Res 176, 268–280. 10.1016/0014-4827(88)90330-8.

van Paassen, B.W., van der Kooi, A.J., van Spaendonck-Zwarts, K.Y., Verhamme, C., Baas, F., and de Visser, M. (2014). PMP22 related neuropathies: Charcot-Marie-Tooth disease type 1A and Hereditary Neuropathy with liability to Pressure Palsies. Orphanet J Rare Dis 9, 38. 10.1186/1750-1172-9-38.

Velapoldi, R.A., and Tonnesen, H.H. (2004). Corrected emission spectra and quantum yields for a series of fluorescent compounds in the visible spectral region. J Fluoresc 14, 465–472. 10.1023/b:jofl.0000031828.96368.c1.

Verhamme, C., King, R.H., ten Asbroek, A.L., Muddle, J.R., Nourallah, M., Wolterman, R., Baas, F., and van Schaik, I.N. (2011). Myelin and axon pathology in a long-term study of PMP22-overexpressing mice. J Neuropathol Exp Neurol 70, 386–398. 10.1097/NEN.0b013e318217eba0.

Vincent, R.K., and Odorico, J.S. (2009). Reduced serum concentration is permissive for increased in vitro endocrine differentiation from murine embryonic stem cells. Differentiation 78, 24–34. 10.1016/j.diff.2009.03.006.

Visigalli, D., Capodivento, G., Basit, A., Fernandez, R., Hamid, Z., Pencova, B., Gemelli, C., Marubbi, D., Pastorino, C., Luoma, A.M., et al. (2020). Exploiting Sphingo- and Glycerophospholipid Impairment to Select Effective Drugs and Biomarkers for CMT1A. Front Neurol 11, 903. 10.3389/fneur.2020.00903.

Walkley, S.U., and Suzuki, K. (2004). Consequences of NPC1 and NPC2 loss of function in mammalian neurons. Biochim Biophys Acta 1685, 48–62. 10.1016/j.bbalip.2004.08.011.

Wang, H., Becuwe, M., Housden, B.E., Chitraju, C., Porras, A.J., Graham, M.M., Liu, X.N., Thiam, A.R., Savage, D.B., Agarwal, A.K., et al. (2016). Seipin is required for converting nascent to mature lipid droplets. Elife 5. 10.7554/eLife.16582.

Yu, C., M. Alterman & R. T. Dobrowsky (2005). Ceramide displaces cholesterol from lipid rafts and decreases the association of the cholesterol binding protein caveolin-1. Journal of Lipid Research, 46, 1678–1691.

Zhou, Y., Miles, J.R., Tavori, H., Lin, M., Khoshbouei, H., Borchelt, D.R., Bazick, H., Landreth, G.E., Lee, S., Fazio, S., and Notterpek, L. (2019). PMP22 Regulates Cholesterol Trafficking and ABCA1-Mediated Cholesterol Efflux. J Neurosci 39, 5404–5418. 10.1523/JNEUROSCI.2942-18.2019.

